# Circulating immune signatures reveal targetable inflammatory pathways in anaplastic thyroid carcinoma

**DOI:** 10.64898/2026.05.19.726015

**Authors:** Pepijn van Houten, Titus Schlüter, Nicholas A Sumpter, Prashant Changoer, Liesbeth van Emst, Leonie S Helder, Julia IP van Heck, Joost HA Martens, Janneke EW Walraven, Petronella B Ottevanger, Han J Bonenkamp, Johannes HW de Wilt, Mihai G Netea, Martin Jaeger, Romana T Netea-Maier

## Abstract

Anaplastic thyroid carcinoma (ATC) is one of the most lethal malignancies. Immune dysregulation is believed to play an important role in ATC. Here, we aimed to characterize the systemic inflammation and the function of circulating immune cells of patients with ATC. First, we retrospectively assessed biochemical parameters of patients with ATC and observed that high systemic inflammation correlated with worse survival. Next, we prospectively investigated the inflammatory proteome, single-cell peripheral blood mononuclear cell transcriptome and epigenetic changes. Circulating concentrations of proinflammatory cytokines were increased in ATC patients. This proinflammatory profile was apparent at the level of gene transcription and chromatin accessibility, especially in monocytes. These findings were substantiated by an increased capacity of peripheral blood mononuclear cells of ATC patients to produce IL-6, IL-8 and lactate. As IL-6 is known to promote tumor cell survival, we assessed its capacity to influence ATC cell proliferation. Blocking IL-6/gp130/Jak/STAT3 pathway inhibited proliferation of ATC cell lines *in vitro*. In conclusion, these findings show that ATC is characterized by inappropriate systemic inflammation and epigenetic and transcriptional reprogramming of circulating monocytes. Proinflammatory cytokines released by monocytes support survival and proliferation of ATC tumor cells, suggesting a therapeutic potential of targeting this pathway in ATC patients.

## Introduction

The large majority of patients with well differentiated follicular derived thyroid carcinoma (DTC) have an excellent prognosis. In contrast, advanced metastatic TC, including the radioiodine refractory TC (RAIR-TC), the poorly differentiated (PDTC), and particularly the undifferentiated (anaplastic) thyroid carcinoma (ATC) tumors pose important therapeutic challenges. ATC is a rare but very aggressive malignancy with a dismal prognosis, characterized by a median survival of less than 6 months (1, 2). For ATC patients, the treatment options are limited and almost never curative. Although new treatment options such as kinase inhibitors and immune checkpoint inhibitors (ICIs) have emerged in recent years for the treatment of cancer (3, 4), most patients with ATC do not respond to multikinase inhibitors or ICIs, and many lack actionable molecular targets for specific kinase inhibitors. Moreover, only a small proportion of treated patients achieve durable responses, hence the urgent need for new treatment options. Furthermore, ICIs are currently not approved for use in for ATC patients (5).

We hypothesize that divergent outcomes observed in patients with differentiated and undifferentiated tumors might be related to distinct pathogenetic mechanisms, particularly those involving tumor-associated inflammatory responses. Unlike DTC, ATCs are heavily infiltrated by myeloid immune cells, and several studies show that tumor-associated macrophages (TAMs) can constitute a substantial proportion of the tumor mass (6–8). It has been also shown that in many malignant tumors TAMs are reprogrammed towards an immunosuppressive phenotype by several tumor-related factors including hypoxia, lactate, and certain cytokines and chemokines (9). These immunosuppressive TAMs promote tumor growth, metastasis, and angiogenesis through secretion of proinflammatory cytokines and growth factors (10). They also suppress anti-tumor immune cells such as CD8^+^ T-lymphocytes, thereby hindering the antitumor efficacy of ICI therapy. Therefore, targeting immunosuppressive myeloid cells is an attractive strategy that may enhance the effectiveness of T-lymphocyte targeting treatments (11).

Paradoxically, cancer-associated immunosuppression often coexists with systemic inflammation, which contributes to clinical manifestations such as fatigue and cachexia, and potentially drives the tumor progression. Only a few studies have investigated immune dysregulation in ATC patients. They reported marked systemic inflammation, including leukocytosis and high concentrations of C-reactive protein (CRP) (12–14). Despite its clear pathophysiological relevance, the inflammatory landscape and the innate immune cells function in patients with ATC remain poorly understood. Moreover, the cellular and molecular mechanisms underlying immune dysregulation in these patients are unknown. Our group has previously shown that in patients with DTC, circulating monocytes and myeloid bone marrow progenitors exhibit transcriptional and functional alterations even before entering the tumor microenvironment (15). Whether similar changes occur in more aggressive TC subtypes, such as ATC, is not yet known. Defining the phenotype and function of myeloid cells in patients with ATC may therefore be essential for identifying new therapeutic targets. Since tumor-promoting inflammation is a hallmarks of cancer and contributes to immunosuppression, understanding how systemic inflammation arises in ATC and how it influences tumor progression is of critical importance (16).

In the present study we aimed to comprehensively characterize systemic inflammation and the phenotype of myeloid cells in ATC patients. To this end, we retrospectively analyzed clinical and biochemical data from a large cohort of patients with ATC and PDTC. In addition, we conducted prospective studies to examine the circulating myeloid cells from ATC patients at the epigenetic, transcriptional, and functional level, and compared these findings with those from other TC subtypes and healthy controls.

## Results

### Baseline characteristics

In total, 119 patients were included in the retrospective study. Of these, 89 patients were diagnosed with ATC and 30 patients with PDTC. Four patients were excluded from this cohort: two with ATC (one because of multiple myeloma and one because of chronic lymphatic leukemia) and two with PDTC (one because of chronic lymphatic leukemia and one because of chronic myeloid leukemia). Baseline characteristics of the two subgroups are shown in Supplementary Table 1. As expected, the median survival from patients with ATC (4 months) was significantly shorter than that of patients with PDTC (35 months).

In total, 107 participants were included in the prospective study. Table 1 shows baseline characteristics of all participants. Healthy controls and DTC patients were younger than patients with RAIR-TC, PDTC or ATC. PDTC and ATC patients were more frequently female compared with the other subgroups. DTC and RAIR-TC subgroups consisted mostly of papillary thyroid carcinoma. Treatment history and metastatic patterns were heterogeneous among the aggressive TC subgroups. Supplementary Figure 1 shows the number of participants from each subgroup that were included per analysis.

**Table 1.** Baseline characteristics of patients in the prospective cohort.

|  | Healthy controls (n=24) | DTC (n=33) | RAIR-TC (n=25) | PDTC (n=10) | ATC (n=15) |
| --- | --- | --- | --- | --- | --- |
| Age (years, median with IQR) | 53 (37-63) | 50 (42-65) | 68 (62-77) | 67 (64-72) | 73 (62-75) |
| Sex (n male (%)) | 12 (50.0%) | 20 (60.6%) | 12 (48.0%) | 2 (20.0%) | 5 (33.3%) |
| DTC subtype | NA |  |  | NA | NA |
| - Papillary TC |  | - 30 | - 17 |  |  |
| - Follicular TC |  | - 0 | - 3 |  |  |
| - Oncocytic TC |  | - 3 | - 5 |  |  |
| Metastases | NA |  |  |  |  |
| - Cervical lymph nodes |  | - 27 | - 7 | - 4 | - 8 |
| - Distant lymph nodes |  | - 0 | - 11 | - 2 | - 6 |
| - Lungs |  | - 1 | - 21 | - 5 | - 6 |
| - Liver |  | - 0 | - 3 | - 0 | - 0 |
| - Bones |  | - 0 | - 8 | - 2 | - 3 |
| - Other |  | - 0 | - 1 | - 3 | - 3 |
| Previous treatment | NA |  |  |  |  |
| - Thyroidectomy |  | - 2 | - 22 | - 6 | - 3 |
| - Radioiodine |  | - 2 | - 22 | - 5 | - 0 |
| - Radiotherapy |  | - 0 | - 7 | - 2 | - 0 |
| - Systemic therapy |  | - 0 | - 2 | - 3 | - 0 |

DTC: differentiated thyroid carcinoma RAIR-TC: radioiodine refractory thyroid carcinoma PDTC: poorly differentiated thyroid carcinoma ATC: anaplastic thyroid carcinoma IQR: interquartile range

### Patients with ATC exhibit systemic inflammation which is associated with a poor prognosis

A comparison of biochemical parameters between ATC and PDTC patients at diagnosis showed significantly higher CRP concentrations, erythrocyte sedimentation rate (ESR) and white blood cell counts in the ATC patients compared with the PDTC patients, suggesting the presence of inappropriate systemic inflammation (Figure 1A). In addition, the neutrophil-lymphocyte-ratio (NLR) was higher at diagnosis in the ATC patients, while the lymphocyte-monocyte-ratio (LMR) was lower than in PDTC patients. This could indicate a shift from lymphopoiesis towards myelopoiesis in the ATC patients, a pattern consistent with systemic inflammation. No differences were observed for albumin concentrations and platelet-lymphocyte ratio (PLR).

**Figure 1.**
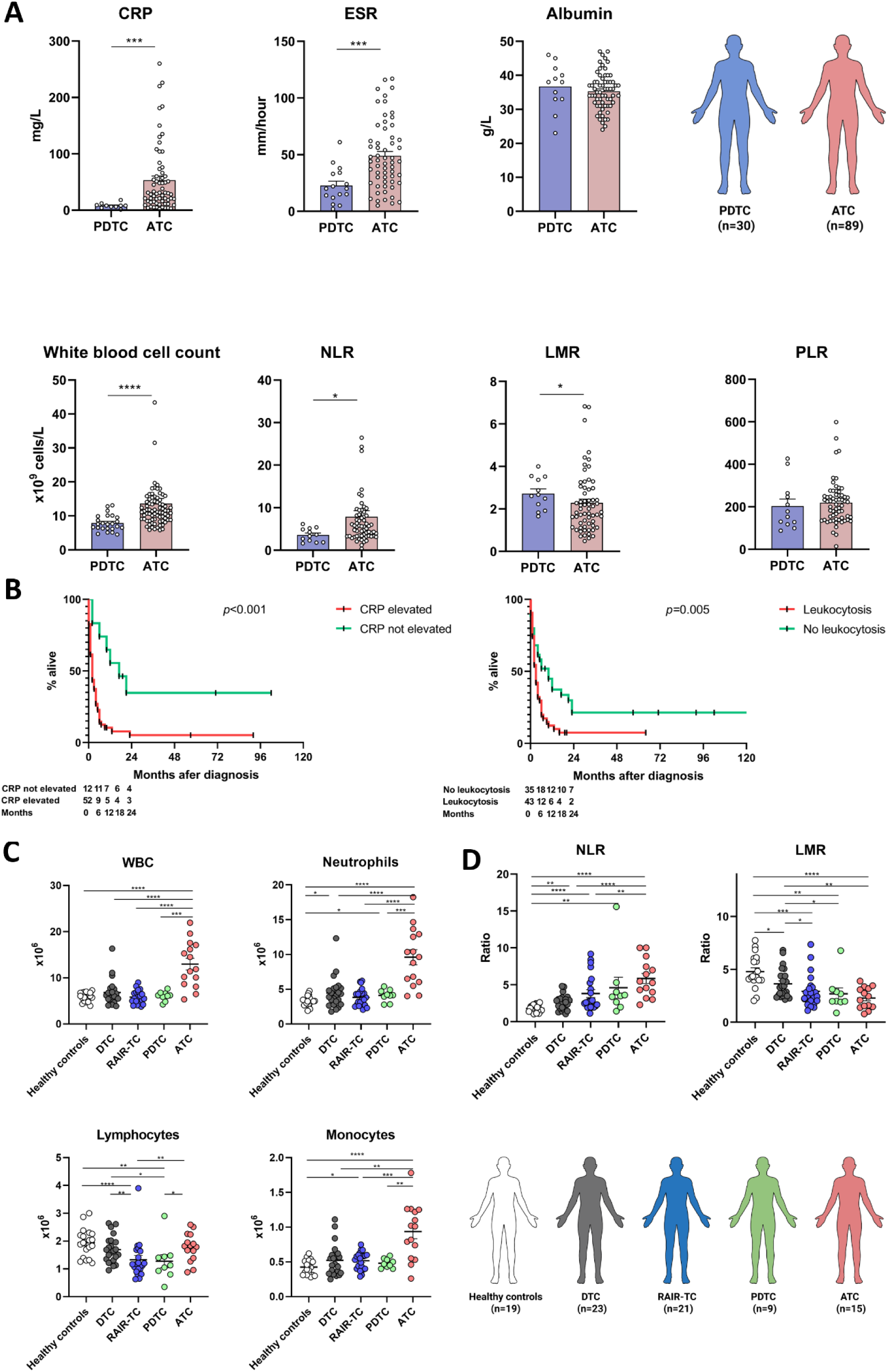
Pronounced systemic inflammation in patients with ATC, correlating negatively with survival. **A:** Comparison of several inflammatory biomarkers in PDTC patients and ATC patients. **B:** Kaplan-Meier curves of ATC patients with elevated CRP at diagnosis compared with ATC patients with normal CRP at diagnosis and ATC patients with leukocytosis at diagnosis compared with ATC patients without leukocytosis at diagnosis. **C:** Absolute numbers of circulating white blood cells, neutrophils, lymphocytes and monocytes in whole blood. **D:** NLR and LMR in whole blood. *: *p*<0.05 **: *p*<0.01 ***: *p*<0.001 ****: *p*<0.0001. ATC: anaplastic thyroid carcinoma. PDTC: poorly differentiated thyroid carcinoma. RAIR-TC: radioiodine-refractory thyroid carcinoma. DTC: differentiated thyroid carcinoma. CRP: C-reactive protein. ESR: Erythrocyte sedimentation rate. NLR: Neutrophil-to-lymphocyte-ratio. LMR: lymphocyte-to-monocyte-ratio. PLR: platelet-to-lymphocyte-ratio.

To identify factors associated with prognosis, a regression analysis was performed. In univariate regression analysis, age at diagnosis, lymph node metastases at diagnosis, distant metastases at diagnosis, hypoalbuminemia, elevated CRP, leukocytosis, and higher NLR all were significantly negatively associated with overall survival (OS) (Supplementary Table 2). There was no significant association between sex, elevated ESR, anemia, LMR, and PLR and OS. In a multivariable model, both age at diagnosis and elevated CRP remained significantly negatively associated with OS. A log-rank test confirmed that, when ATC patients with elevated and normal CRP concentrations were compared, the survival was significantly worse for the patients with elevated CRP at diagnosis. The same finding was observed comparing ATC patients with or without leukocytosis at diagnosis (Kaplan-Meier curves are shown in Figure 1B).

### Increased white blood cell counts, particularly neutrophils and monocytes in patients with ATC

In the prospective cohort, white blood cell (WBC) counts were higher in patients with ATC compared with the other subgroups, mostly due to higher numbers of neutrophils (Figure 1C). Neutrophil numbers were also increased in patients with DTC or PDTC compared with healthy controls, while lymphocyte numbers were lower in patients with RAIR-TC and PDTC. Furthermore, monocyte counts were higher in ATC patients compared with all other subgroups.

When assessed as percentages of WBC instead of absolute numbers, neutrophils were present in higher proportions in the more aggressive phenotypes ATC and PDTC, compared with the three other subgroups (Supplementary Figure 2). For lymphocyte percentages, the opposite was observed, with lower lymphocyte percentages in ATC and PDTC patients. For monocyte percentages, less obvious differences were observed, with only higher percentages in RAIR-TC patients compared with healthy controls or ATC patients.

NLR was higher in all TC subgroups compared with healthy controls (Figure 1D). NLR was also higher in ATC patients compared with DTC and RAIR-TC patients, but not with PDTC patients. Conversely, LMR was lower in all TC subgroups compared with healthy controls (Figure 1D). In addition, LMR was higher in DTC patients compared with the three more aggressive TC subgroups (ATC, PDTC and RAIR-TC).

### ATC patients display a distinct plasma proteome with increased concentrations of IL-6 family cytokines and growth factors

We subsequently assessed the systemic inflammatory proteome by a proximity extension assay that measured 92 inflammation-related proteins. The plasma proteome was analyzed for a subset of 23 DTC patients, 17 RAIR-TC patients, 10 PDTC patients and 10 ATC patients. Of the 92 proteins in the panel, 83 proteins were measurable in at least 75% of the samples and were included in the analyses. The proteome of the ATC subgroup was clearly distinct from that of the other subgroups (PCA plot in Figure 2A). Comparing the ATC to the DTC subgroup, the concentrations of 37 proteins were upregulated, while the concentrations of two proteins (TRANCE and SCF) were lower (Figure 2B). Concentrations of nineteen proteins were increased when comparing the ATC to the RAIR-TC subgroup (these all were also increased when comparing ATC to DTC), while three were decreased (TRANCE, SCF and Flt3-ligand) (Figure 2C). Comparing ATC to PDTC, only three proteins were upregulated (interleukin-6 (IL-6), oncostatin-M (OSM) and CCL23), while two proteins were downregulated (SCF and Flt3-ligand) (Figure 2D). IL-6, OSM and CCL23 were elevated in comparisons between ATC and all other three subgroups.

**Figure 2.**
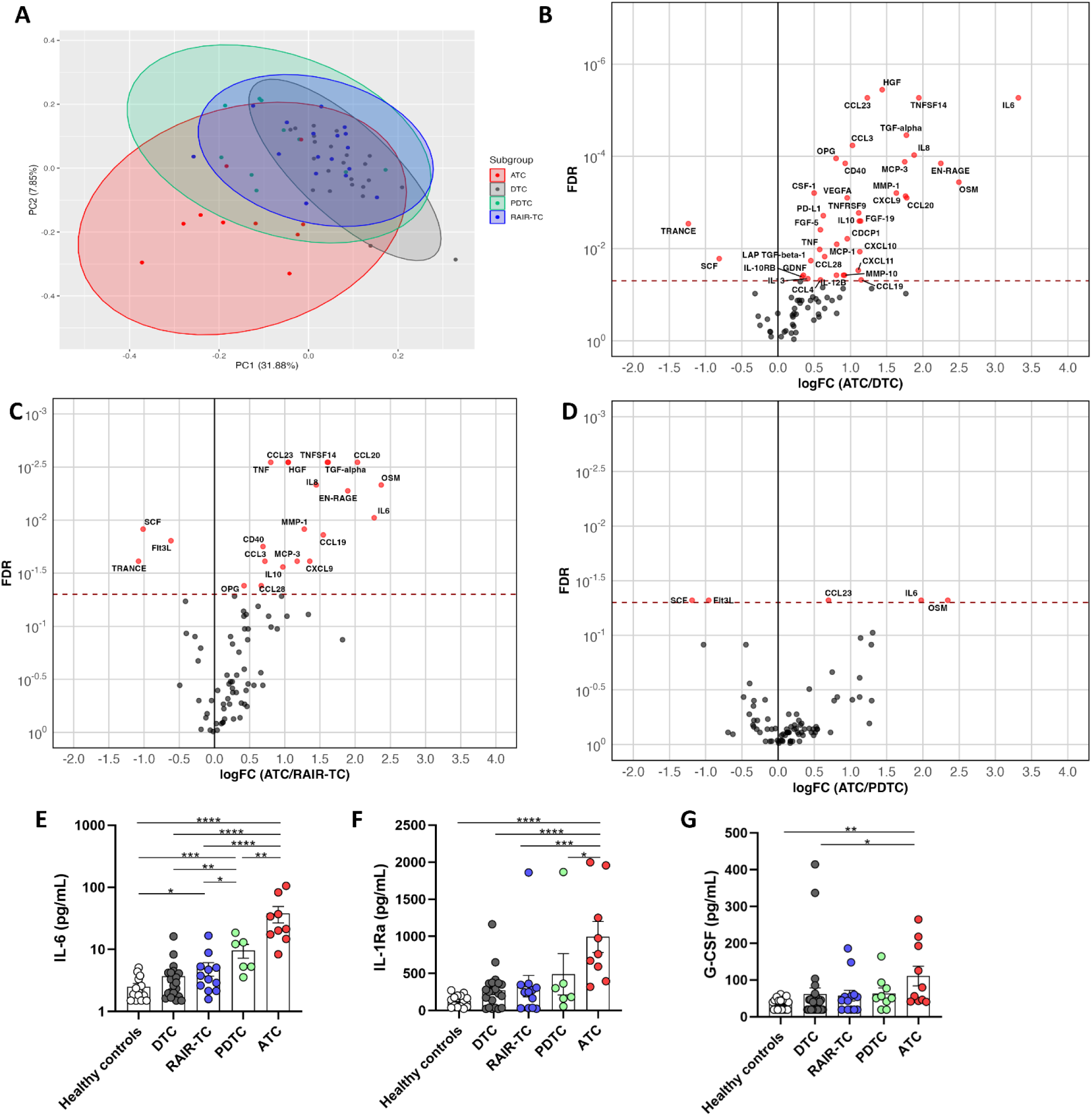
Upregulation of inflammatory biomarkers in the circulation of patients with ATC. **A:** Principal component analysis of inflammatory plasma proteome of different subgroups. B-D: Vulcano plots showing differentially expressed proteins in plasma from ATC patients compared with DTC **(B)**, RAIR-TC **(C)** and PDTC **(D)**. E-G: Concentrations of circulating IL-6 **(E)**, IL-1Ra **(F)** and G-CSF **(G)** in plasma. *: *p*<0.05 **: *p*<0.01 ***: *p*<0.001 ****: *p*<0.0001. ATC: anaplastic thyroid carcinoma. PDTC: poorly differentiated thyroid carcinoma. RAIR-TC: radioiodine-refractory thyroid carcinoma. DTC: differentiated thyroid carcinoma. G-CSF: granulocyte colony stimulating factor.

Compared to DTC, PDTC patients showed an increased concentration of 31 proteins and no decreased proteins in the circulation (Supplementary Figure 3A). Of these 31 proteins, 22 were also increased in the ATC/DTC comparison. Comparison of PDTC and RAIR-TC subgroups demonstrated no differences in protein concentration (Supplementary Figure 3B) and comparison of RAIR-TC to DTC showed only two increased proteins (Flt3-ligand and CX3CL1 (Supplementary Figure 3C).

To provide an independent validation of the plasma proteome findings, the concentrations of several circulating cytokines and growth factors were measured by enzyme-linked immunosorbent assay (ELISA). IL-6 plasma concentrations showed a gradient from lowest concentrations in healthy controls to highest concentrations in ATC patients (Figure 2E). Likewise, IL-1-receptor antagonist (IL-1Ra) concentrations were significantly higher in plasma from ATC patients than in plasma from all other subgroups (Figure 2F). IL-1β concentrations were undetectable in 64% of samples (46/72) (Supplementary Figure 4A), and were not different between subgroups. Circulating concentrations of both osteopontin and CXCL9 were highest in patients with ATC and PDTC (Supplementary Figure 4B & 4C). No differences were observed for osteopontin or CXCL9 concentrations between healthy controls, DTC patients and RAIR-TC patients.

To assess the role of growth factors that have stimulatory effects on myelopoiesis, granulocyte colony-stimulating factor (G-CSF) and (granulocyte-macrophage colony-stimulating factor (GM-CSF) were measured in plasma. G-CSF plasma concentration was highest in plasma from ATC patients (Figure 2G) and showed significantly higher concentrations when compared with those found in healthy controls and DTC patients. GM-CSF was only detectable in samples from three donors (one DTC patient and two ATC patients, Supplementary Figure 4D).

### Transcriptional changes in circulating monocytes from ATC patients

Immune cells in the circulation play a central role in host defense and pathophysiology of many diseases, including cancer. To characterize the transcriptional phenotype of circulating immune cells, single-cell RNA-sequencing was performed on PBMCs from five patients with ATC, five patients with DTC and four healthy controls (Figure 3A). In total, 34,043 cells were sequenced. We identified several known subpopulations within the PBMCs, based on marker expression (Supplementary Figure 5A). Importantly, after annotating differentially expressed (DE) genes per cell type, the highest DE count was observed in CD14^+^ monocytes (Figure 3B), arguing for an important role of monocyte function dysregulation in ATC.

**Figure 3.**
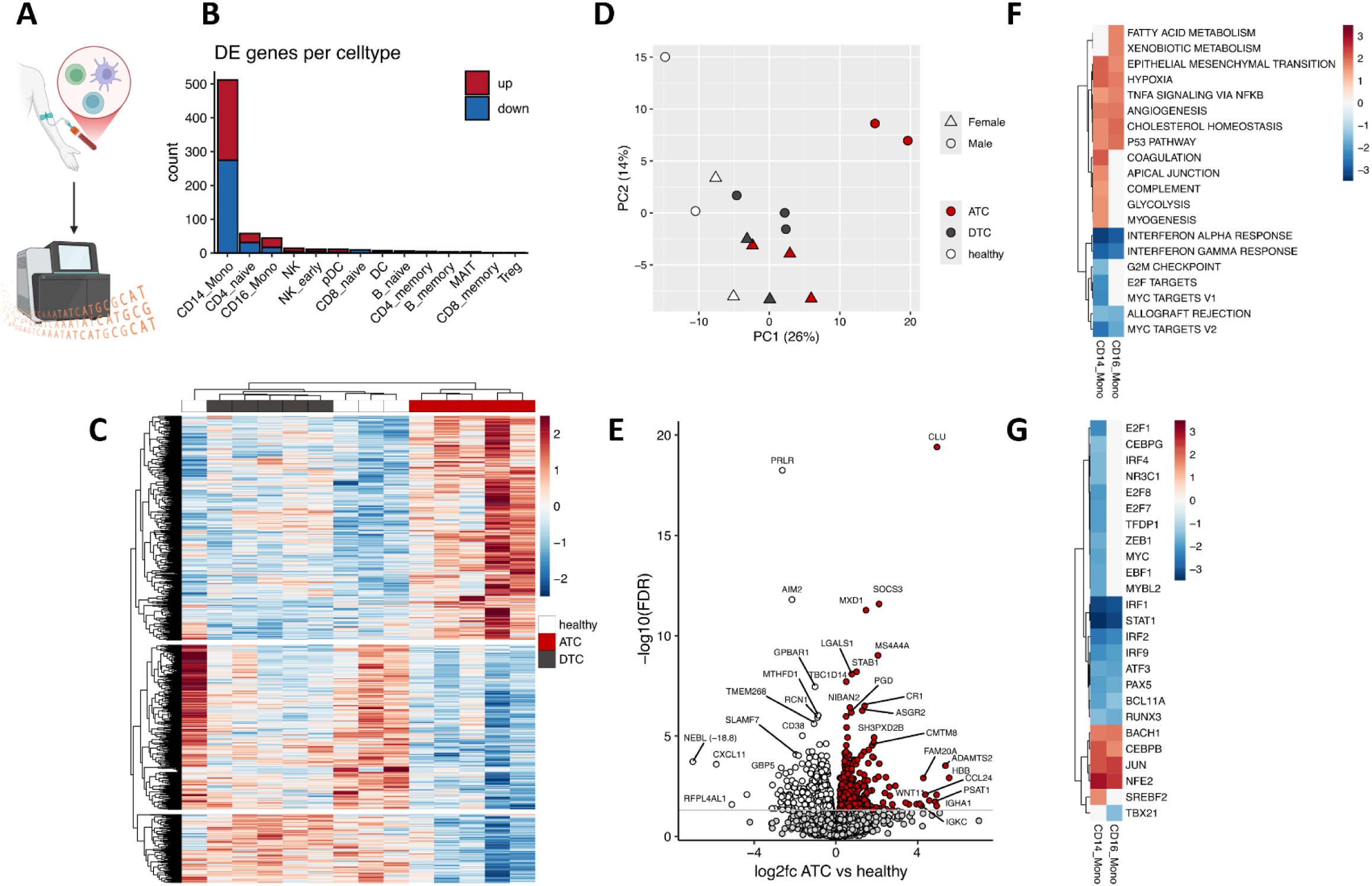
Transcriptomic analysis of monocytes from ATC patients showes upregulation of proinflammatory and protumoral pathways and downregulation of interferon related pathways. **A:** Single-cell RNA sequencing was performed on isolated PBMCs. **B:** Differentially expressed genes per cell type, comparing ATC patients with healthy controls. **C:** Heatmap of differentially expressed genes in CD14+ monocytes showing clustering of ATC patients. **D:** Principal component analysis of CD14+ monocytes. **E:** Volcano plot showing up- and downregulated genes in CD14+ monocytes of ATC patients compared with healthy controls. **F:** Significantly up- and downregulated pathways in monocytes from ATC patients compared with healthy controls in monocytes. Colours represent normalized enrichement scores. **G:** Significantly up- and downregulated regulons in monocytes from ATC patients compared with healthy controls. Colours represent normalized enrichment scores. DE: differentially expressed, GSEA: gene set enrichment analysis, ATC: anaplastic thyroid carcinoma, DTC: differentiated thyroid carcinoma.

Comparison between monocyte transcriptional programs in ATC patients, DTC patients and healthy controls, showed distinct patterns for the three subgroups (heatmap in Figure 3C and PCA plot in Figure 3D). Significantly up- and downregulated genes are shown in the Volcano plots comparing ATC patients to healthy controls (Figure 3E) and DTC patients (Supplementary Figure 5B). Pathway analysis of the DEGs in monocytes from ATC patients showed upregulation of several pathways important for carcinogenesis, including Glycolysis, Hypoxia, Epithelial-mesenchymal transition and Angiogenesis (Figure 3F and Supplementary Figure 6A-B). Conversely, gene sets corresponding to type I and type II interferon pathways were the most significantly downregulated in monocytes of ATC patients (Supplementary Figure 6C). Analysis of transcription factor expression showed that several interferon regulatory factors (IRF1, IRF2, IRF4 and IRF9), E2Fs, MYC, and STAT1 were downregulated in monocytes of ATC patients compared with healthy controls (Figure 3G and Supplementary Figure 6D-E). The upregulated transcription factors in monocytes from ATC patients included JUN, NFE2, and CEBPB.

### Monocytes from ATC patients depict epigenetic rewiring of inflammation and interferon pathways

As monocytes of ATC patients represented the immune cell population with most changes in the transcriptome, we investigated whether these changes were due to modifications of epigenetic control of gene transcription. Chromatin accessibility was assessed by ATAC-sequencing on circulating monocytes from eight patients with ATC (six female and two male, median age: 65.5 years) and eight healthy controls (five female and three male, median age: 63.5 years) (Figure 4A). After correction for sex, distinct epigenetic profiles could be observed for both groups, as depicted in the PCA analysis (Figure 4B). In total, 1,039 differentially accessible regions were identified (Figure 4C). Next, we performed transcription factor motif analysis using these regions. The ten known motifs with the highest similarity for the up- and downregulated motifs are shown in Supplementary Tables 3 & 4. We found strong enrichment of the Jun motif, which is involved in transcription factor AP-1 (Figure 4D). Conversely, the downregulated peaks in the monocytes from ATC patients were involved in the interferon-stimulated response element (ISRE) motif (Figure 4E). ISRE is a promotor sequence that acts as a binding site for interferon stimulated transcription factors, such as IRFs. ISRE was the most enriched motif based on the downregulated peaks in monocytes from ATC patients, in line with the concept that type I and type II IFN pathways are the most downregulated in the RNA-sequencing analysis of monocytes from ATC patients.

**Figure 4.**
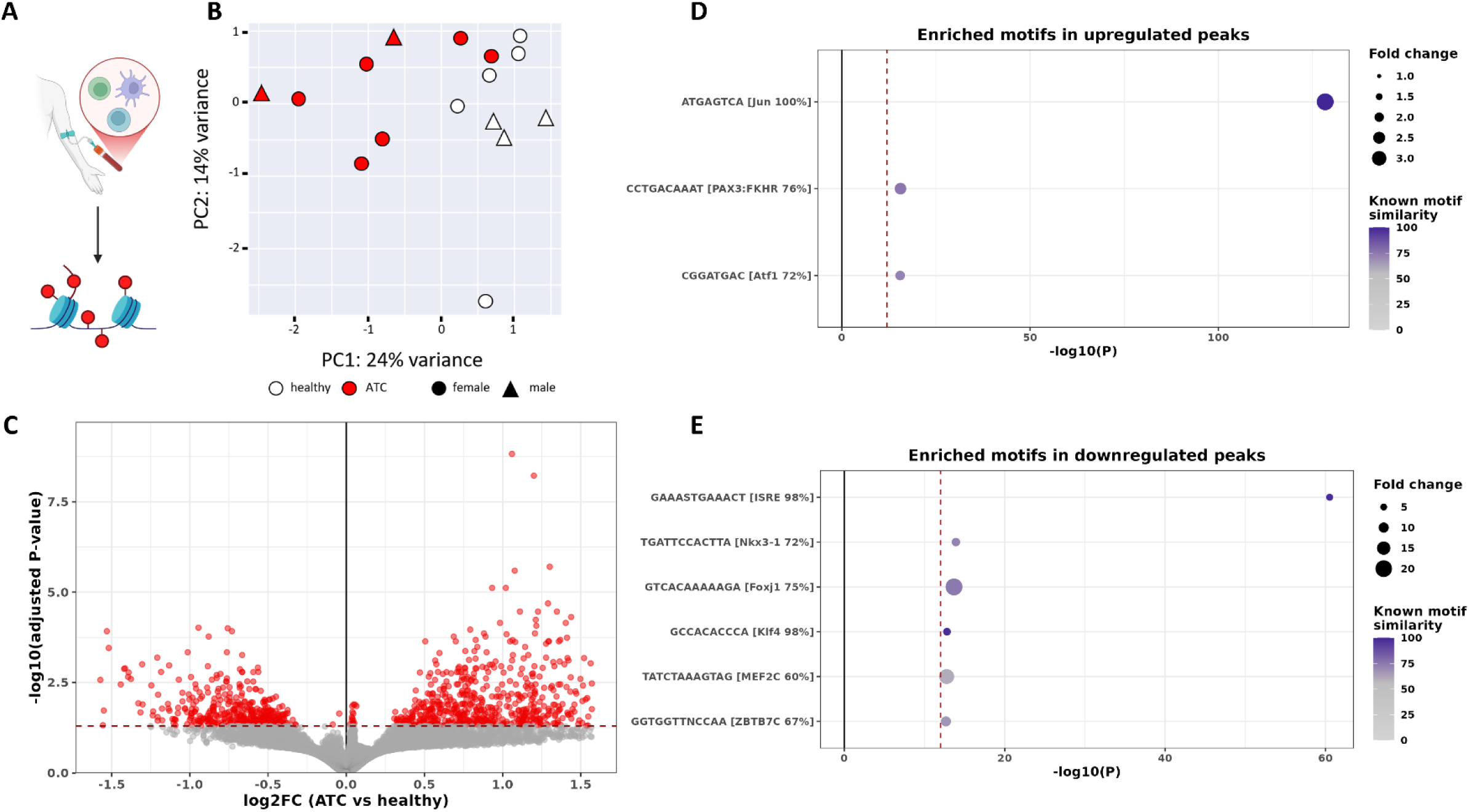
Distinct epigenetic rewiring of monocytes from patients with ATC, with upregulation of AP-1 and downregulation of ISRE. **A:** Bulk ATAC-sequencing was performed on isolated monocytes. **B:** Principal component analysis of ATAC-sequencing results for healthy controls and anaplastic thyroid cancer patients, corrected for sex. **C:** Volcano plot of significantly differentially accessible peaks in ATC patients compared with healthy controls. **D:** Significantly enriched motifs based on the upregulated peaks in monocytes from ATC patients. **E:** Significantly enriched motifs based on the downregulated peaks from ATC patients. ATC: anaplastic thyroid carcinoma; ISRE: interferon-sensitive response element.

### PBMCs from ATC patients show increased IL-6 and IL-8, and reduced IFN-γ production capacity

We then aimed to functionally validate the hypothesis that myeloid cells from ATC patients have a different profile in the induction of inflammatory pathways compared to healthy controls and patients with other TC subtypes. Monocyte-derived cytokine and lactate production capacity were assessed by stimulating PBMCs for 24 hours with LPS, P3C, IL-1α or *C. albicans* (Figure 5A). In line with the observed transcriptional and epigenetic changes in monocytes from ATC patients, as well as the elevated concentrations of proinflammatory cytokines in circulation, the production of IL-6 by PBMCs from ATC patients was significantly higher compared with healthy controls and DTC patients (Figure 5B and Supplementary Figure 7A). In addition, PBMCs from patients with PDTC and RAIR-TC produced more IL-6 compared with healthy controls. Similarly, IL-8 production by PBMCs from ATC patients was significantly higher compared with healthy controls and DTC patients for all tested stimuli (Figure 5C and Supplementary Figure 7B). For P3C and *C. albicans* stimulations, PBMCs from RAIR-TC patients produced more IL-8 compared with healthy controls. Furthermore, IL-1Ra was more strongly produced upon stimulation of PBMCs from all TC subtypes compared with healthy controls, with the highest concentrations being produced by PBMCs from TC patients with the more aggressive subtypes (Figure 5D and Supplementary Figure 7C). Conversely, production of IL-1β was not increased in stimulated PBMCs from ATC patients, compared with healthy controls or other TC subtypes (Supplementary Figure 8A). There was a significantly increased production of IL-1β for DTC patients (P3C and *C. albicans* stimulations) and RAIR-TC patients (LPS and P3C stimulations) compared with healthy controls. No differences were observed in the IL-10 production by PBMCs from the different subgroups, except for higher production of IL-10 after stimulation by P3C of PBMCs of DTC patients compared with healthy controls (Supplementary Figure 8B). Lactate production was increased in LPS-stimulated PBMCs from ATC patients compared with PBMCs from healthy controls, DTC patients and RAIR-TC patients (Figure 5E), in line with the known importance of glycolysis for the inflammatory function of monocytes (17).

**Figure 5.**
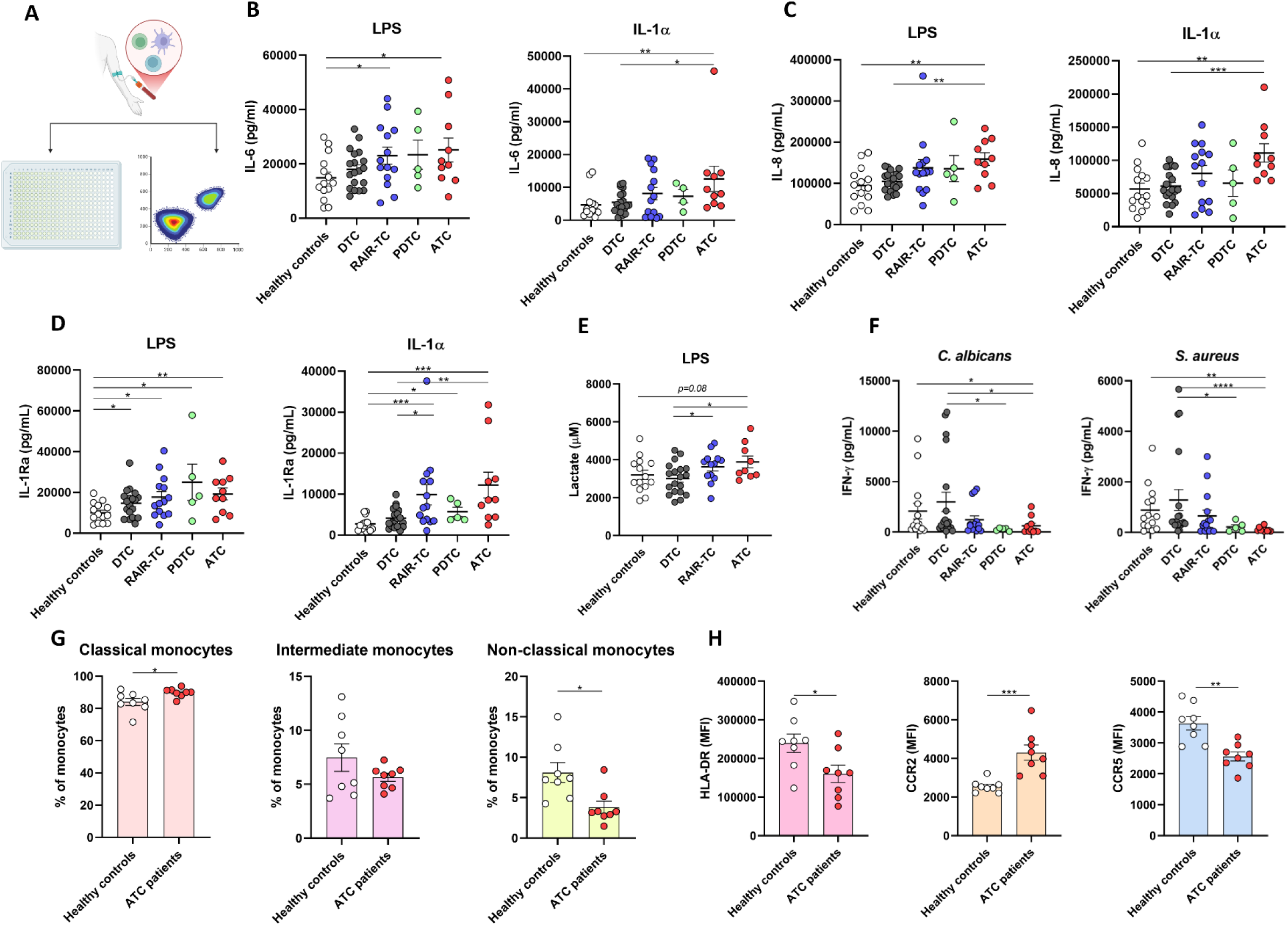
Monocytes from patients with ATC display increased cytokine production capacity. **A:** ELISA was performed on stimulated PBMCs and flowcytometry was performed on isolated monocytes. Production of IL-6 **(B)**, IL-8 **(C)** and IL-1Ra **(D)** by monocytes after 24 hours of stimulation by LPS or IL-1α. **E:** Production of lactate by monocytes after 24 hours of stimulation by LPS. **F:** Production of IFN-γ after 7 days of stimulation by *Candida albicans* or *Staphylococcus aureus*. **G:** Percentages of subpopulations of monocytes as part of all circulating monocytes. **H:** Expression of markers HLA-DR, CCR2 and CCR5 on monocytes. *: *p*<0.05 **: *p*<0.01 ***: *p*<0.001 ****: *p*<0.0001. ATC: anaplastic thyroid carcinoma; PDTC: poorly differentiated thyroid carcinoma; RAIR-TC: radioiodine refractory thyroid carcinoma; DTC: differentiated thyroid carcinoma.

To assess lymphocyte-derived cytokine production, PBMCs were stimulated for 7 days with *C. albicans* or *S. aureus*. Upon both stimulations, production of interferon-γ (IFN-γ) was lower in PBMCs from patients with ATC compared with healthy controls and patients with DTC (Figure 5F). In addition, PBMCs from PDTC patients also produced low amounts of IFN-γ. In contrast, IL-17 and IL-22 production were similar for the different subgroups upon stimulation (Supplementary Figure 9).

### Higher proportion of classical monocytes with decreased HLA-DR and CCR5 expression and increased CCR2 expression in ATC patients

Next, circulating monocytes from eight ATC patients and eight healthy controls underwent immunophenotyping by flow cytometry. Subsets of monocytes were defined as classical, intermediate or non-classical monocytes, based on CD14 and CD16 expression (see gating strategy in Supplementary Figure 10). The monocyte compartment of the ATC patients consisted of relatively more classical monocytes and less non-classical monocytes compared with that of the healthy controls (Figure 5G). Expression of HLA-DR and CCR5 were decreased on monocytes from ATC patients compared with the healthy controls (Figure 5G). Conversely, expression of CCR2 was increased on monocytes from ATC patients (Figure 5H). Expression of HLA-DR, CCR2 and CCR5 per monocyte subpopulation is shown in Supplementary Figure 11. Within the subpopulations of classical monocytes and intermediate monocytes, the median expression of CCR2 was higher in patients with ATC compared with healthy controls.

### Blocking IL-6/JAK/STAT3 pathway results in potent inhibition of ATC cell proliferation

IL-6 and OSM are known as survival factors for malignant cells (18, 19). As shown in the current work, both proteins are more strongly produced by immune cells of ATC patients. Therefore, we subsequently assessed their effect on ATC cell survival and proliferation. ATC cell lines 8505C, CAL62, and 8305C were treated with different concentrations of IL-6 or OSM for 48 hours. Neither IL-6 nor OSM influenced ATC cell proliferation, although there was a significantly higher rate of proliferation after incubation of cell line CAL62 with 1ng/mL of OSM (Supplementary Figure 12).

Subsequently, the same cell lines were treated with inhibitors of different components of the IL-6/JAK/STAT3 pathway: tocilizumab as IL-6-receptor inhibitor, bazedoxifene acetate as gp130 inhibitor (a component of the IL-6 receptor), ruxolitinib as JAK inhibitor, and WP1066 as STAT3 inhibitor (Figure 6A). To exclude cytotoxicity, a lactate dehydrogenase (LDH) assay was performed with the 3 highest concentrations of each compound. The high concentrations of bazedoxifene acetate, and to a lesser extent also WP1066, resulted in high levels of LDH, and were thus cytotoxic (Supplementary Figure 14). When assessing cell proliferation based on ATP production, treatment with tocilizumab inhibited cell proliferation only in 8505C and 8305C cell lines at high concentrations (Figure 6B), and bazedoxifene acetate inhibited cell proliferation at 10 µM, and to a lesser extent at 2.5 µM (Figure 6C). Ruxolitinib and WP1066 were potent inhibitors of ATC cell proliferation, with the same patterns in all different cell lines (Figures 6D-E).

**Figure 6.**
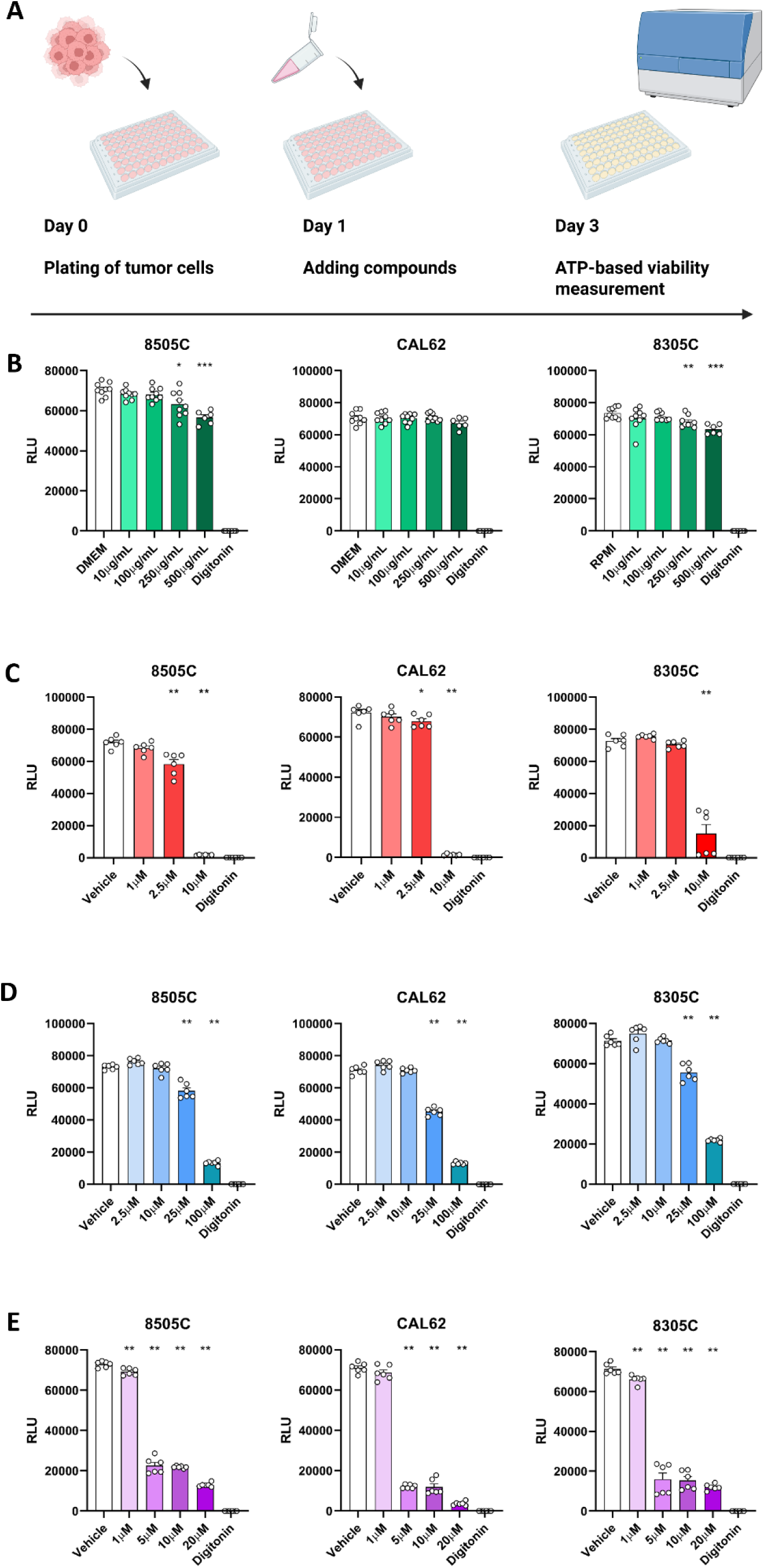
Blocking the IL-6/gp130/Jak/STAT3 pathway inhibits proliferation of ATC cells *in vitro*. **A:** Overview of experiments with ATC cell lines and inhibitors of IL-6/gp130/Jak/STAT3 pathway. Viability is measured after 48 hours of treatment, using an ATP-based assay. B-E: Viability of ATC cell lines 8505C, CAL62 and 8305C after 48 hours of treatment with IL-6 receptor antagonist tocilizumab **(B)**, gp130 inhibitor bazedoxifene acetate **(C)**, Jak inhibitor ruxolitinib **(D)** and STAT3 inhibitor WP1066 **(E)**. *: p<0.05 **: p<0.01 ***: p<0.001

## Discussion

The present study provides the first comprehensive evaluation of the innate immune landscape and systemic inflammatory profile in patients with ATC. We report that ATC is marked by sustained systemic inflammation, which is associated with a poor outcome. This inflammatory state is accompanied by extensive epigenetic and transcriptional rewiring of circulating monocytes in ATC patients, resulting in two major functional consequences: an increased production and release of proinflammatory cytokines that provide survival signals to malignant cells, and an impaired interferon signaling pathway, essential for activation of cytotoxic T-cells. Importantly, blockade of IL-6 family-mediated inflammatory signaling inhibited proliferation and survival of ATC tumor cell lines, highlighting this pathway as a promising therapeutic target.

ATC tumors are known to be heavily infiltrated by immunosuppressive and protumoral myeloid cells (6–8), suggesting that myeloid cell dysregulation contributes substantially to disease aggressiveness. Our study support this hypothesis by demonstrating a profoundly dysregulated systemic inflammatory phenotype in ATC, in which myeloid cells are likely central players. First, we observed markedly elevated concentrations of systemic inflammatory biomarkers in circulation, which negatively correlated with survival. Second, epigenetic, transcriptional, and functional analyses revealed that circulating monocytes from ATC patients exhibited pronounced proinflammatory and proangiogenic phenotypes compared with monocytes from patients with less aggressive TC subtypes and healthy controls. Third, PBMCs from patients with ATC displayed reduced IFN-γ production capacity, accompanied by reduced chromatin accessibility at IFN-related loci. Finally, pharmacological inhibition of the gp130/Jak/STAT3-axis, which can be activated by members of the IL-6 family, significantly reduced ATC cells proliferation. Taken together, these findings strongly implicate myeloid cells dysregulation as a key contributor to ATC pathogenesis and supports an approach targeting inflammation as a future treatment strategy in ATC.

Using both retrospective and prospective cohorts, we demonstrated elevated systemic inflammatory biomarkers in patients with ATC, such as increased neutrophil counts, CRP and IL-6 concentrations. Tumor-promoting inflammation is a recognized hallmark of cancer progression (16). Inflammation facilitates several protumoral processes, including angiogenesis, epithelial-to-mesenchymal transition, and metastasis. In our retrospective cohort, elevated CRP, leukocytosis, and high NLR at diagnosis were associated with a poor survival in ATC. This is in line with a recent multicenter cohort study in ATC patients where high NLR at baseline was associated with worse survival (20). The high prevalence of neutrophilia and monocytosis in ATC patients likely reflects emergency myelopoiesis, by increased production of growth factors such as G-CSF and cytokines such as IL-1 and IL-6, resulting in increased differentiation of bone marrow progenitors into myeloid cells and more potent mobilization of these myeloid cells into the circulation (21, 22). Indeed circulating concentrations of G-CSF and IL-6 were indeed significantly increased in our prospective ATC cohort compared with other TC subgroups and healthy controls.

Proteomic profiling further substantiated the inflammatory phenotype of ATC. Plasma from ATC patients contained elevated concentrations of several circulating biomarkers including proangiogenic proteins, chemokines, matrix metalloproteinases and various interleukins compared with other TC subtypes. Notably, IL-6 and OSM, both members of the IL-6 family, and the chemokine CCL23, were strongly elevated in ATC patients in comparison to every other TC subtype, including the PDTC. In contrast, minimal differences were observed between the inflammatory proteomes of DTC and RAIR-TC patients, despite distinct clinical outcome, suggesting that additional non-inflammatory mechanisms contribute to disease progression in these differentiated tumors.

Single-cell RNA-sequencing and ATAC-sequencing revealed that the dysregulation of inflammation in ATC patients was accompanied by important changes in the transcriptome and chromatin accessibility profiles of their circulating monocytes. The circulating monocytes from patients with ATC exhibited the highest number of differentially expressed genes (DEGs) among circulating immune populations as compared to control individuals. Mirroring the increase in circulating inflammatory biomarkers, several genes and pathways involved in cytokine production were upregulated in monocytes from ATC patients, with Jun (a transcription factor that in combination with Fos forms AP-1), being strongly upregulated at both transcriptomic and epigenetic levels. Functional assays confirmed enhanced production of IL-6, IL-8, and lactate by monocytes from ATC patients. AP-1 is known to be involved in regulation of the IL-6 and IL-8 genes, and could partially explain the pronounced proinflammatory signature that was observed in the ATC patients (23, 24). Given its important role in regulating several protumoral processes including cell growth, angiogenesis and drug resistance, AP-1 has been proposed as potential target for new cancer treatments (25). Considering also the rich myeloid infiltrate in ATC tumors, highly believed to be responsible for the lymphocyte suppression in the tumor microenvironment, the present data strongly position the monocyte-driven immune dysregulation as a key node in ATC-associated inflammation.

Conversely, type 1 IFNs and IFN-γ pathways were downregulated in monocytes from patients with ATC. The reduced expression of several IRFs and STAT1 and strong downregulation of the ISRE motif accessibility suggests impaired IFN responsiveness. IFNs are known to have acute antineoplastic effects on cancer cells, and can stimulate immune cells including NK cells, T cells and dendritic cells towards anticancer functions (26–28). Type 1 IFNs can induce an antitumoral phenotype in tumor-associated neutrophils (29) and is long known to have antitumoral effects (30). Selective intratumoral activation of type 1 IFNs has shown promising results in preclinical models with synergy to ICIs (31, 32). The antitumoral effects of IFN-γ (33), and its capacity to upregulate PD-L1 expression on epithelial cells (34), could be leveraged for by combining IFN-γ with ICI treatment for synergistic effects, as was recently shown in a phase I trial in patients with various solid tumors (35). Therefore, therapeutic strategies aimed at restoring IFN signaling, potentially in combination with ICIs, warrant further investigation in ATC.

Phenotypic analysis further demonstrated a shift towards classical monocytes and a lower proportion of non-classical monocytes in ATC patients as compared with healthy controls. Non-classical monocytes are involved in several antitumoral processes including the recruitment of NK cells and CD8^+^ T-cells, and inhibition of regulatory T-cells (36–38) whereas circulating classical monocytes serve as major source of protumoral TAMs (39). Additionaly, the decreased expression of HLA-DR on monocytes from ATC patients, compared with healthy controls, suggests an immunosuppressive phenotype similar to CD14^+^HLA-DR^lo^ monocytes described in patients with other malignancies such as pancreatic cancer and breast cancer, and shown to negatively correlate with survival in patients with renal cell carcinoma (40–42). Targeting and reprogramming this monocyte subpopulation represents a potential therapeutic avenue to be explored in future studies.

IL-6 emerged as a central mediator across multiple levels of analysis. Both systemic circulating concentrations and monocyte-derived secretion of IL-6 were significantly increased in ATC patients compared with healthy controls and other TC subtypes. IL-6 is produced by several cell types including tumor cells, fibroblasts and immune cells such as monocytes and neutrophils. Together with OSM, which was also elevated in the plasma of ATC patients, IL-6 signals through the gp130/Jak/STAT3-axis to promote tumor cell proliferation, survival, metastasis and angiogenesis (18). Pharmacological inhibition of the gp130/Jak/STAT3-axis at different levels, including Inhibition of Jak (ruxolitinib) and STAT3 (WP1066), reduced proliferation of several ATC cell lines *in vitro*, consistent with prior preclinical *in vitro* data in other cancer types (43–45). Guo *et al*. showed recently that administration of ruxolitinib in a mouse xenograft model with ATC cell line 8505C resulted in decreased tumor growth (43). phase I trial showed acceptable safety of WP1066 in patients with glioblastoma, a tumor type with high levels of p-STAT3, similar to ATC (46). A phase II trial with WP1066 is currently being initiated (NCT05879250). These findings support further clinical evaluation of this pathway in ATC, including its effects on tumor-associated myeloid cells.

Our study also has limitations. Retrospective data may inherently be subject to information bias, although this is unlikely to have affected immunological assays. Functional analyses primarily focused on myeloid cells, as the most important cell population in inflammatory processes. Nonetheless, future studies should extend investigation of additional immune cell populations. Moreover, the cohort consisted exclusively of patients of European ancestry, and validation in more diverse populations is warranted. Nevertheless, there are also important strengths of the study. This is the first comprehensive assessment of innate immunity and inflammation in ATC patients, and the relation with their clinical outcome. In addition, our study include multi-omics profiling of the immune function at different levels (ATAC-sequencing, RNA-sequencing, proteomics assays), integrated with robust functional validation and clinical data.

In conclusion, ATC is characterized by high levels of tumor-associated systemic inflammation and extensive, epigenetic and transcriptional rewiring of myeloid cells, resulting in a strong functional proinflammatory bias, coupled with impaired IFN pathway activity. This dual immune dysregulation likely contributes to defective lymphocyte activation, angiogenesis and tumor progression. Our findings highlight myeloid cells and IL-6 family signaling as central drivers of ATC pathogenesis and identify these pathways as promising therapeutic targets. More broadly, this work provides a framework for investigating innate immune dysregulation across other malignancies as well.

## Methods

### Retrospective cohort study

Medical records of patients treated for ATC or PDTC at the Radboud University Medical Center, Nijmegen, the Netherlands, between 1999 and March 2025 were examined retrospectively. Patients with inflammatory or infectious comorbidities and patients treated with immunomodulating medication were excluded from the analysis. Data were collected on demographic characteristics, diagnostic procedures, tumor stage at diagnosis, treatment, and survival. In addition, results from biochemical analysis at diagnosis were collected, more specifically: albumin, CRP, ESR, WBC, neutrophil count, lymphocyte count, monocyte count, and platelet count. The following ratios were calculated: NLR, LMR, and PLR. Biochemical parameters were considered as elevated when the value was higher than the upper limit of normal at the time of the assay.

### Prospective cohort study

Patients aged ≥ 18 years old who were managed at the Radboud University Medical Center for DTC, advanced and metastatic RAIR-TC, PDTC, or ATC were included in this study between September 2022 and April 2025. All patients had structural disease at inclusion. Exclusion criteria were: inflammatory or infectious comorbidities, use of medication that interfered with the immune system, other active malignancies, pregnancy, or a self-reported alcohol consumption of >21 units per week. The study was approved by the Research Ethics Committee (2022-16025). Healthy controls for experiments were included via a separate study “Blood donation for experimental *in vitro* research” which was approved by Medical Ethical Committee Oost-Nederland (2023-16366). All study procedures were performed in accordance with the Principles of the Declaration of Helsinki.

### Materials

Blood was drawn using ethylenediaminetetraacetic acid (EDTA) blood collection tubes, after written informed consent was obtained. EDTA-plasma was obtained from whole blood by centrifugation and stored at -80 °C until further use. All experiments were performed in Roswell Park Memorial Institute (RPMI) 1640 Dutch Modified medium (Gibco), supplemented with gentamycin 50 µg/mL, pyruvate 1 mM and glutamax 2 mM. The used stimuli are described in the Supplementary Materials.

### Cell counts

Cell counts were obtained in whole blood using a Sysmex automated hematology analyzer (XN-450; Sysmex Corporation).

### Olink targeted proteomics

Using the Inflammation Target 96 panel from Olink proteomics, 92 inflammation related proteins were measured in EDTA plasma (47). All samples were measured in one batch. Four internal controls and two external controls were included in the assay. The analysis of the proteomics results is described in the Supplementary Materials.

### Single-cell RNA-sequencing

PBMC samples were frozen in CryoStor CS10 (StemCell Technologies) and stored at -150° C. Upon use, PBMCs were thawed at 37° C and diluted with 14 mL of RPMI + 10% heat-inactivated Fetal Bovine Serum (FBS-26A, Capricorn) containing 12.5 ug/mL DNase I (740963, Macherey Nagel).

PBMCs were fixed and frozen din 50% glycerol until further processing (10x Genomics demonstrated protocol CG000478). Multiplex libraries containing a barcode for each donor were prepared following the Chromium Fixed RNA Profiling user guide for multiplexed samples and capturing 2,000 cells per sample (CG000527). Libraries were sequenced on a NovaSeqX 10B with 100 cycles at 20,000 reads per cell. The analyses of sequencing results are described in the Supplementary Materials.

### ATAC-sequencing

Samples for ATAC-sequencing were prepared from MACS-sorted monocytes. 5.0x10^4^ cells were washed in cold PBS and resuspended in a transposase reaction mix (12.5 µL 2x Tagment DNA buffer, 10.25 µL H_2_O, 2 µL Tagment DNA Enzyme (Illumina) and 0.25 µL digitonin (Promega). Samples were incubated for 30 minutes at 37 °C and then purified using the MinElute kit (QIAGEN) according to the manufacturer’s guidelines. The eluted 10 µL was stored at -20 °C until further processing. Processing and analysis are described in the Supplementary Materials.

### PBMC stimulation

PBMCs were isolated from whole blood using Ficoll-Paque PLUS (GE Healthcare) in SepMate isolation tubes (STEMCELL Technologies). After isolation, PBMCs were suspended in RPMI medium and counted. For *ex vivo* stimulation, 0.5x10^6^ PBMCs were plated in duplicate in 96-well round-bottom plates. PBMCs were stimulated either for 24 hours with LPS (10 ng/mL), P3C (10 µg/mL), IL-1α (10 ng/mL), *C. albicans* (1x10^6^/mL) or RPMI as negative control or for 7 days with *C. albicans* (1x10^6^/mL), *S. aureus* (1x10^6^/mL) or RPMI as negative control. The medium of the 7 days stimulation experiment was supplemented with 10% human pooled serum. After stimulation, plates were centrifuged and supernatants were collected and stored at -20 °C until analysis. Procedures for the measurement of cytokines and lactate are described in the Supplementary Materials.

### Flow cytometry

5.0x10^5^ monocytes were stained with Fixable Viability Stain 620 (BD Biosciences) in 100 µL of PBS. After staining, cells were washed with PBA (PBS + 1% bovine serum albumin + 2 mM EDTA) and centrifuged. Cells were resuspended in PBA with 10% TruStain FcX Receptor Blocker (Biolegend). After incubation, cells were stained for 15 minutes at room temperature with the antibody panel: CD14-FITC (Biolegend) HLA-DR-PerCP/Cyanine5.5 (Biolegend), CCR2-BV421 (BD Biosciences), CD16-BV510 (Biolegend) & CCR5-BV650 (BD Biosciences). Antibodies were diluted in PBA with 10% Brilliant Stain buffer (BD Biosciences). After the final incubation, cells were washed with PBA, centrifuged and resuspended in 200 µL PBA. Stained samples and unstained controls were then measured on a CytoFLEX flow cytometer (Beckman Coulter).

### ATC cell line experiments

ATC cell lines 8505C and CAL62 were cultured in Dulbecco’s Modified Eagle Medium (DMEM) and 8305C was cultured in RPMI 1640 Dutch modification (both Gibco). Both culture media were supplemented with penicillin-streptomycin 100 U/mL (Gibco), glutamax 2 mM (Gibco) and 10% Fetal Calf Serum (Capricorn Scientific).

At day 0, 50 µL of medium containing 4.0x10^3^ (8305C) or 2.0x10^3^ cells (8505C and CAL62) were plated on 96-well flat bottom plates in triplicate and incubated overnight. On day 1, the following compounds were added: tocilizumab (Tyenne®, Fresenius Kabi, 1-500 µg/mL), bazedoxifene acetate (Sanbio, 1-25 µM), ruxolitinib (Sanbio, 2.5-100 µM), WP1066 (Merck, 1-20 µM), recombinant IL-6 (Miltenyi Biotec, 10-250 ng/mL) or recombinant OSM, (R&D Bio-Techne, 1-100 ng/mL). For DMSO dissolved compounds, a DMSO control was added (culture medium + 0.1% DMSO). Culture medium supplemented with digitonin (Promega, 200 µg/mL) was added as a positive control. After 48 hours, cell viability was measured using the ATP-based CellTiter-Glo® Luminescent Cell Viability Assay (Promega) according to the manufacturer’s protocol.

For the LDH assay, the same procedure was followed. After 48 hours of stimulation, the supernatant was collected. LDH was measured in the supernatant using the Cytotox96® Non-Radioactive Cytotoxicity Assay (Promega) according to the manufacturer’s protocol.

### Statistical analysis

For the retrospective cohort study, baseline characteristics are shown as part of the group or as median with interquartile range (IQR). Characteristics are compared between ATC and PDTC patients by the Chi-square test or Mann-Whitney U test. OS was compared between ATC patients in whom a marker was elevated and ATC patients in whom that marker was not elevated, using the log-rank test. Univariable and multivariable analyses were compared using the Cox proportional hazards model. For the multivariable analysis, 15 events were needed per included parameter. Analyses were performed in SPSS software version 29. *P*<0.05 was considered statistically significant.

For the prospective *ex vivo* study, baseline characteristics are shown as part of the group or as median with IQR. Characteristics are compared between subgroups by the Chi-square test or Mann-Whitney U test. Cell counts and cytokine concentrations are compared between by the Mann-Whitney U test. Flow cytometry data were analyzed in FlowJo software version 10.8.1. Median Fluorescence Index (MFI) of markers of interest were compared between subgroups by the Mann-Whitney U test. Analyses were performed in GraphPad Prism version 8.0.2. Analyses were performed two-tailed and *p*<0.05 was considered statistically significant.

For the LDH assay, all optical density (OD) values were subtracted by the OD values of the negative controls (either medium or DMSO control, depending on the stimulus). Then, the OD values of different conditions were calculated as percentage of the OD value from the cell lysis condition.

### Sex as a biological variable

Our study examined samples of both male and female donors. Sex was regarded as a biological variable in the regression analysis of the retrospective cohort.

## Data availability

All data associated with this study are in the paper or in the Supplementary Materials. Single-cell RNA-sequencing and ATAC-sequencing data are stored and accessible as data access collection on the Radboud Data Repository with collection identifier ru.rumc.tcsarc_t0000594a_dac_655.

## Author contributions

Pvh, MGN, MJ and RTN-M conceived the study plan. PvH, PC, TS, LvE, LSH and MJ performed the experiments. PvH, TS, NAS, JIPvH and JHAM performed the analyses. PvH, TS, NAS and JIPvH visualized the data. PvH and RTN-M wrote the manuscript. All authors reviewed and edited the manuscript. MGN, MJ and RTN-M supervised the study. PvH managed the project administration.

