## Supplementary Materials for "Circulating immune signatures reveal targetable inflammatory pathways in anaplastic thyroid carcinoma"

**Tables**

**Supplementary Table 1**. Baseline characteristics of patients in the retrospective cohort.

|  | ATC (n=89) | PDTC (n=30) | *p-*value |
| --- | --- | --- | --- |
| Sex (*n* male (%)) | 36 (40.4%) | 16 (53.3%) | 0.219 |
| Age at diagnosis (years, median with IQR) | 69 (59-75) | 66 (62-72) | 0.405 |
| Distant metastases at diagnosis (*n* (%)) | 51 (57.3%) | 12 (40.0%) | 0.101 |
| Cervical lymph node metastases at diagnosis (*n* (%)) | 61 (68.5%) | 19 (63.3%) | 0.439 |
| Survival (months, median ± SEM) | 4 ± 3 | 35 ± 12 | <0.001 |
| *Treatment* | | | |
| Best supportive care only (*n* (%)) | 29 (32.6%) | 1 (3.3%) | 0.001 |
| Surgery of primary tumor (*n* (%)) | 39 (43.8%) | 26 (86.7%) | <0.001 |
| Radiotherapy (*n* (%)) | 45 (50.6%) | 14 (46.7%) | 0.712 |
| Chemotherapy (*n* (%)) | 5 (5.6%) | 1 (3.3%) | 0.526 |
| TKI (*n* (%)) | 16 (18.0%) | 11 (36.7%) | 0.035 |
| Immunotherapy (*n* (%)) | 4 (4.5%) | 1 (3.3%) | 0.628 |

ATC: anaplastic thyroid carcinoma PDTC: poorly differentiated thyroid carcinoma IQR: interquartile range TKI: tyrosine kinase inhibitor SEM: standard error of the mean.

**Supplementary Table 2.** Hazard ratios for death in relation to clinical and biochemical parameters at diagnosis in 89 patients with anaplastic thyroid carcinoma.

|  | Univariate regression | | Multivariate regression | |
| --- | --- | --- | --- | --- |
|  | HR (95%-CI) | *p-*value | HR (95%-CI) | *p-*value |
| Age at diagnosis (years) | 1.026 (1.004-1.049) | *0.021* | 1.031 (1.005-1.058) | *0.021* |
| Sex (male versus female) | 1.082 (0.677-1.730) | 0.742 |  |  |
| Lymph node metastases at diagnosis | 3.730 (1.965-7.082) | *<0.001* |  |  |
| Distant metastases at diagnosis | 2.679 (1.600-4.486) | *<0.001* | 1.345 (0.717-2.525) | 0.356 |
| Hypoalbuminemia | 1.893 (1.086-3.300) | *0.024* |  |  |
| Elevated CRP | 3.362 (1.483-7.624) | 0.004 | 2.697 (1.148-6.332) | *0.023* |
| Elevated ESR | 1.916 (0.806-4.559) | 0.141 |  |  |
| Leukocytosis | 1.947 (1.159-3.272) | *0.012* | 1.631 (0.874-3.046) | 0.125 |
| Anemia | 1.514 (0.906-2.530) | 0.113 |  |  |
| NLR | 1.022 (1.004-1.041) | *0.015* |  |  |
| LMR | 1.001 (0.997-1.006) | 0.592 |  |  |
| PLR | 1.002 (0.999-1.005) | 0.136 |  |  |

HR: hazard ratio CRP: C-reactive protein ESR: erythrocyte sedimentation rate NLR: neutrophil-lymphocyte ratio LMR: lymphocyte-monocyte ratio PLR: platelet-lymphocyte ratio

**Supplementary Table 3.** The top ten motifs with the highest known motif similarity for the three significantly upregulated motifs in monocytes from ATC patients, with corresponding motif similarity scores.

| **Motif 1** | | **Motif 2** | | **Motif 3** | |
| --- | --- | --- | --- | --- | --- |
| Jun | 100% | PAX2:FKHR | 76% | Atf1 | 72% |
| FOS | 99% | MEIS1 | 73% | FOSL1::JUND | 70% |
| FOSL1 | 98% | Tgif1 | 72% | MIES1 | 70% |
| BNC2 | 98% | Tgif2 | 68% | JUN::JUNB | 69% |
| AP-1 | 98% | Msgn1 | 68% | MEIS3 | 67% |
| BATF | 97% | MEIS2 | 68% | RHOFXF1 | 67% |
| BATF::JUN | 97% | BCL11A | 67% | Atf1 | 67% |
| FOS::JUN | 97% | Meis1 | 65% | Crx | 67% |
| Atf3 | 97% | Meis2 | 65% | Dmbx1 | 67% |
| FOSL2::JUN | 97% | NF1 | 65% | NR2C1 | 66% |

**Supplementary Table 4.** The top ten motifs with the highest known motif similarity for the six significantly downregulated motifs in monocytes from ATC patients, with corresponding motif similarity scores.

| **Motif 1** | | **Motif 2** | | **Motif 3** | |
| --- | --- | --- | --- | --- | --- |
| ISRE | 98% | Nkx3-1 | 72% | Foxj1 | 75% |
| IRF1 | 98% | ISL2 | 68% | CDx2 | 65% |
| IRF3 | 94% | Nkx3-2 | 67% | Npas4 | 64% |
| STAT1:STAT2 | 93% | Nkx2-3 | 65% | SOX15 | 63% |
| IRF2 | 93% | Hdx | 63% | HOXD10 | 63% |
| IRF8 | 93% | Nanog | 63% | CDX4 | 62% |
| IRF1 | 93% | Nkx3-1 | 63% | HOXC10 | 62% |
| IRF7 | 89% | PU.1:IRF8 | 61% | HOXC9 | 62% |
| IRF2 | 86% | Nkx2-6 | 61% | Cdx1 | 62% |
| PU.1:IRF8 | 86% | RELB | 60% | CDX1 | 62% |
| **Motif 4** | | **Motif 5** | | **Motif 6** | |
| Klf4 | 98% | MEF2C | 60% | ZBTB7C | 67% |
| EKLF | 94% | Mef2a | 60% | ZNF692 | 67% |
| Klf9 | 93% | POU2F2 | 60% | ZNF354C | 67% |
| Klf5 | 90% | POU1F1 | 59% | Zbtb7b | 64% |
| Klf6 | 89% | Mef2c | 59% | PRDM10 | 63% |
| Klf9 | 88% | ZKSCAN1 | 58% | RUNX1 | 61% |
| Klf1 | 94% | GATA3 | 57% | ZBTC7B | 60% |
| Klf3 | 84% | TRPS1 | 56% | Rfx3 | 60% |
| Klf5 | 82% | LIN54 | 56% | Bcl11B | 60% |
| Klf6 | 81% | Mef2b | 56% | ZBTB9 | 60% |

**Figures**

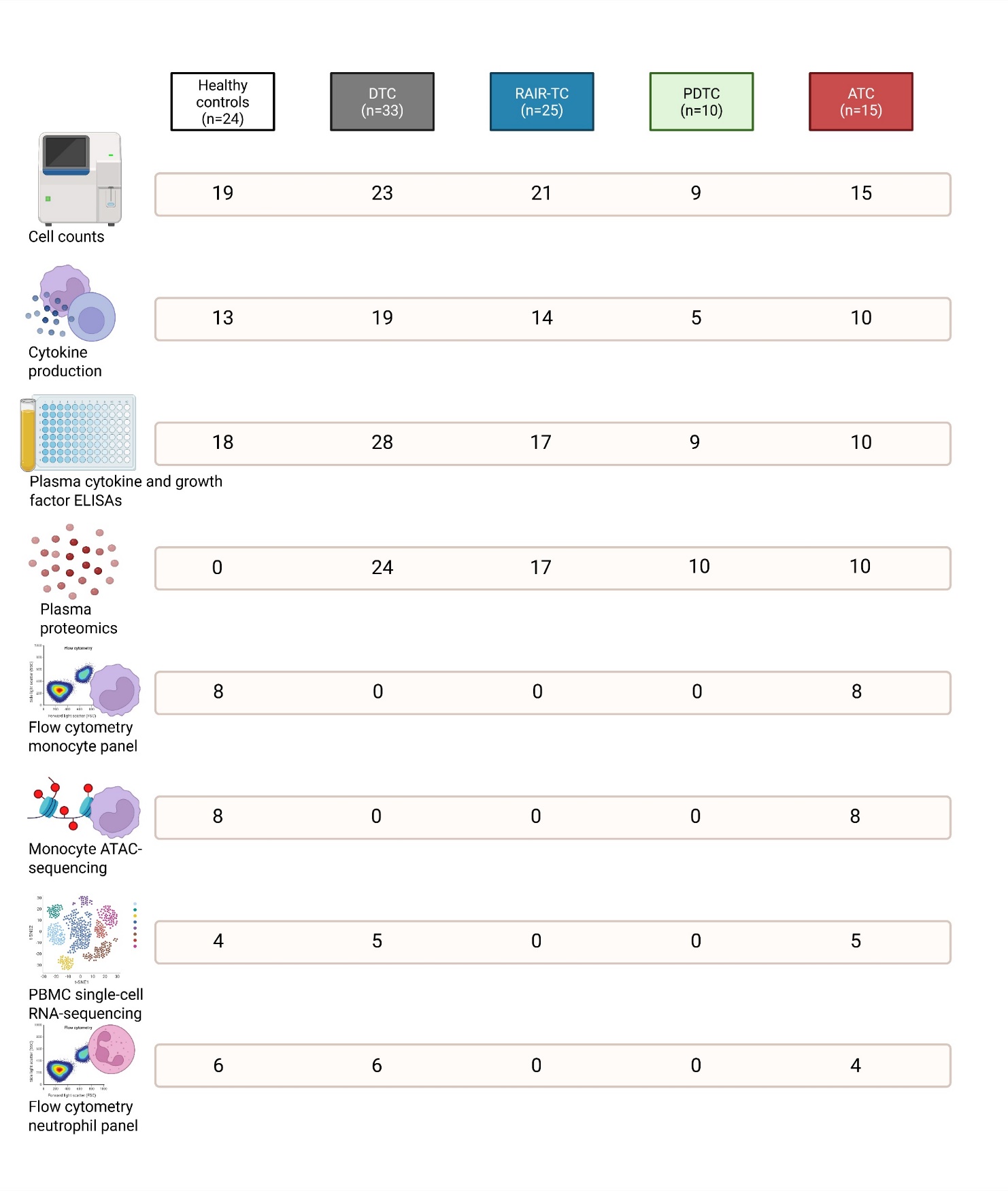
**Supplementary Figure 1.** Assays performed per subgroup of participants.

DTC: Differentiated thyroid carcinoma; RAIR-TC: Radioiodine refractory thyroid carcinoma; PDTC: Poorly differentiated thyroid carcinoma; ATC: anaplastic thyroid carcinoma

**Supplementary Figure 2.** Percentages of neutrophils, lymphocytes and monocytes as part of white blood cells in whole blood.

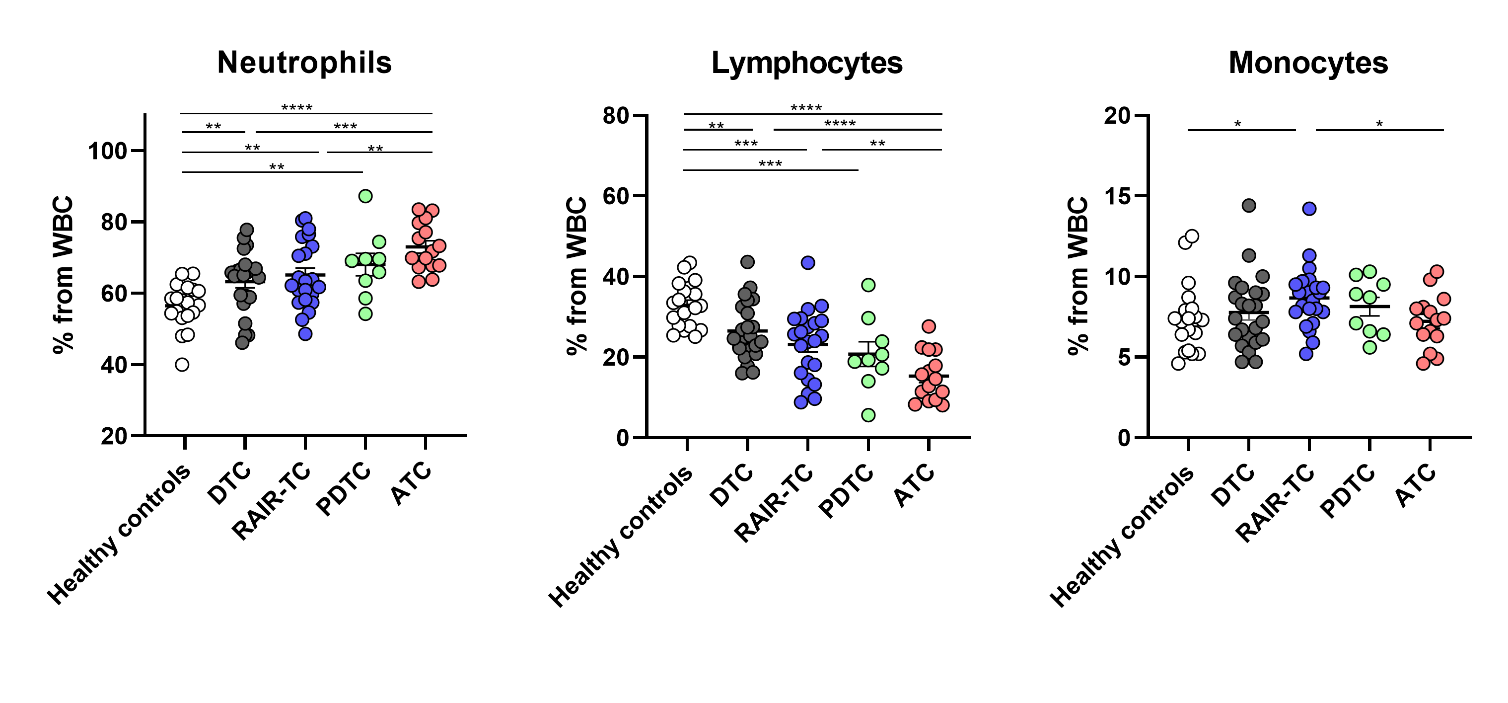

*: *p*<0.05 **: *p*<0.01 ***: *p*<0.001 ****: *p*<0.0001. WBC: white blood cells; DTC: differentiated thyroid carcinoma; RAIR-TC: radioiodine refractory thyroid carcinoma; PDTC: poorly differentiated thyroid carcinoma; ATC: anaplastic thyroid carcinoma.

**Supplementary Figure 3.** Volcano plots showing differentially expressed proteins measured by Olink in plasma. **A:** PDTC compared to DTC. **B:** PDTC compared to RAIR-DTC. **C:** RAIR-DTC compared to DTC.

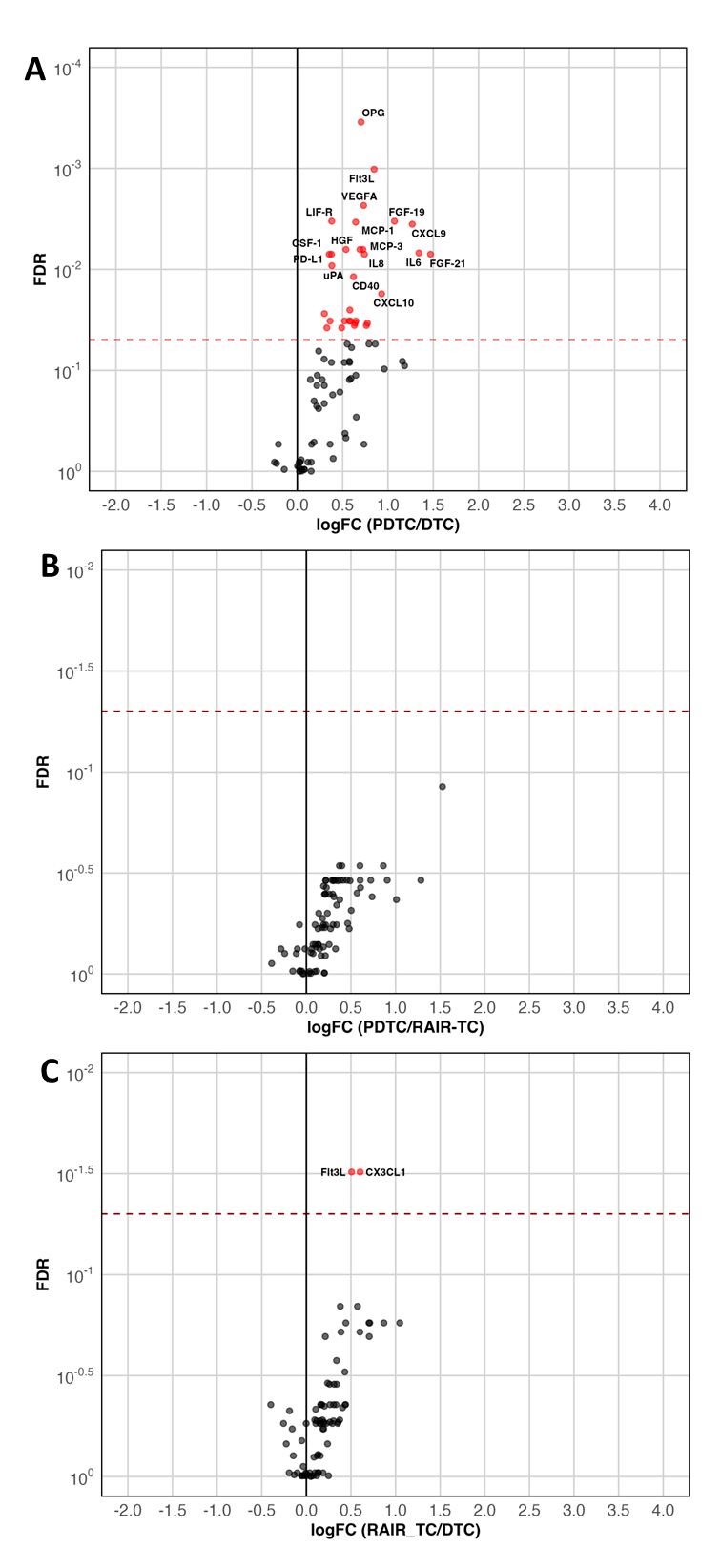

**Supplementary Figure 4.** Plasma concentrations of IL-1β **(A)**, Osteopontin **(B)**, CXCL9 **(C)** and GM-CSF **(D)**.

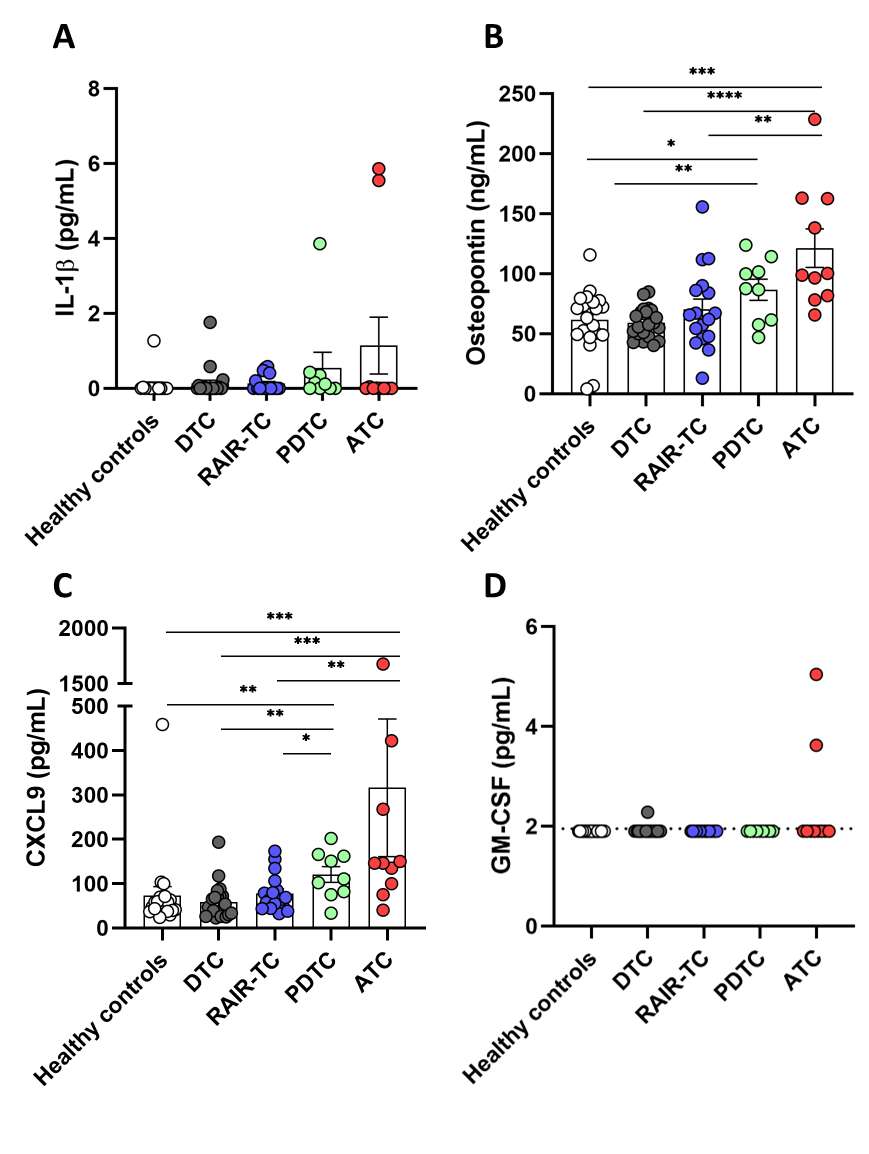

*: *p*<0.05 **: *p*<0.01 ***: *p*<0.001 ****: *p*<0.0001. DTC: differentiated thyroid carcinoma; RAIR-TC: radioiodine refractory thyroid carcinoma; PDTC: poorly differentiated thyroid carcinoma; ATC: anaplastic thyroid carcinoma.

**Supplementary Figure 5. A:** UMAP of single-cell RNA sequencing data in peripheral blood mononuclear cells. **B:** Volcano plot showing differentially expressed genes in CD14+ monocytes of ATC patients compared with DTC patients

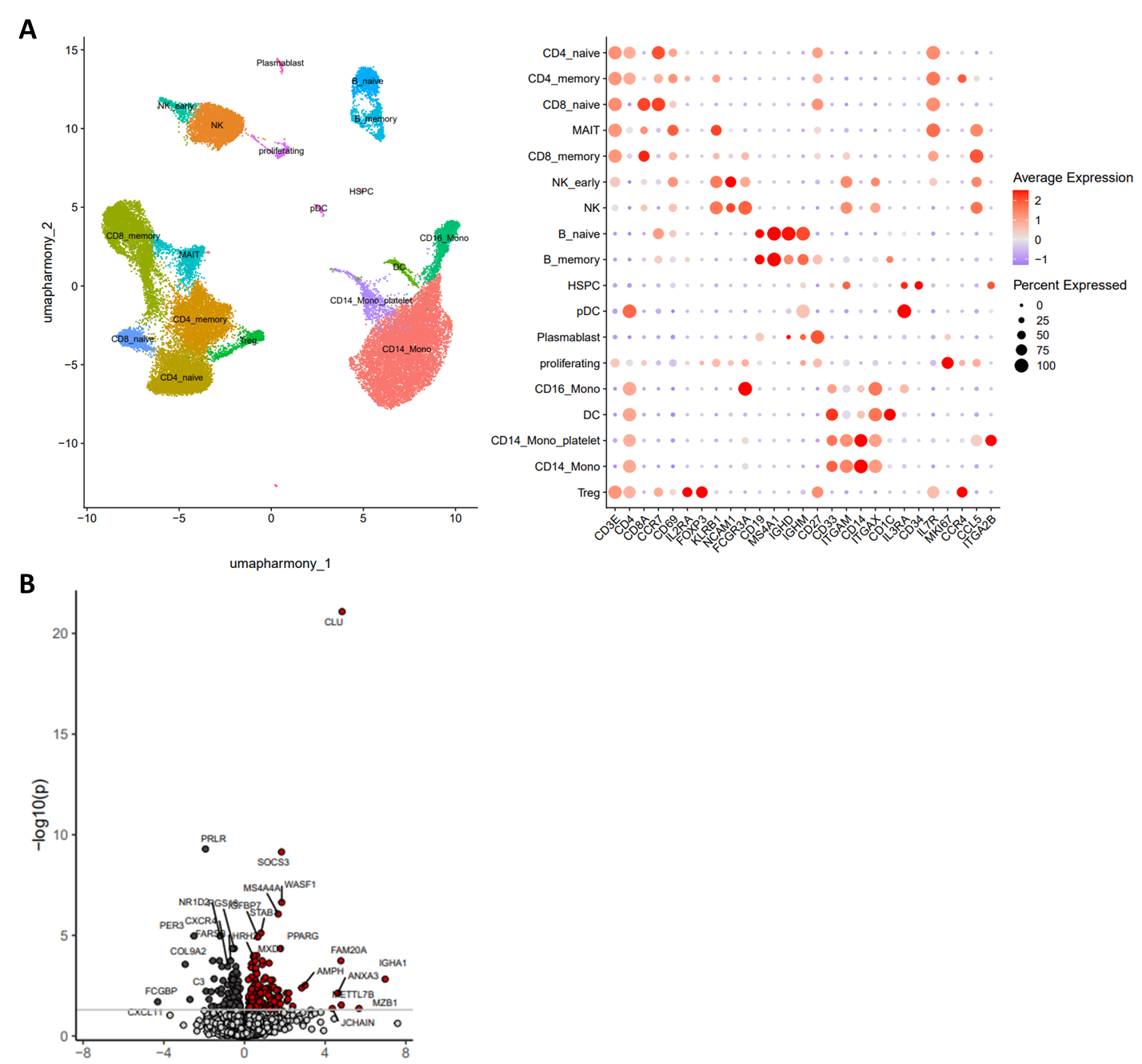

**Supplementary Figure 6.** **A:** Significantly up- and downregulated pathways in ATC patients compared with healthy controls. **B:** Significantly up- and downregulated pathways in ATC patients compared with DTC patients. **C:** Volcano plot showing up- and downregulated pathways in CD14+ monocytes from ATC patients compared with healthy controls. Pathways in black are significant in both GSEA and overrepresentation analysis, pathways in grey are only significant in GSEA. **D:** Significantly up- and downregulated regulons in ATC patients compared with healthy controls. **E:** Significantly up- and downregulated regulons in ATC patients compared with DTC patients. Legends represent normalized enrichment scores.

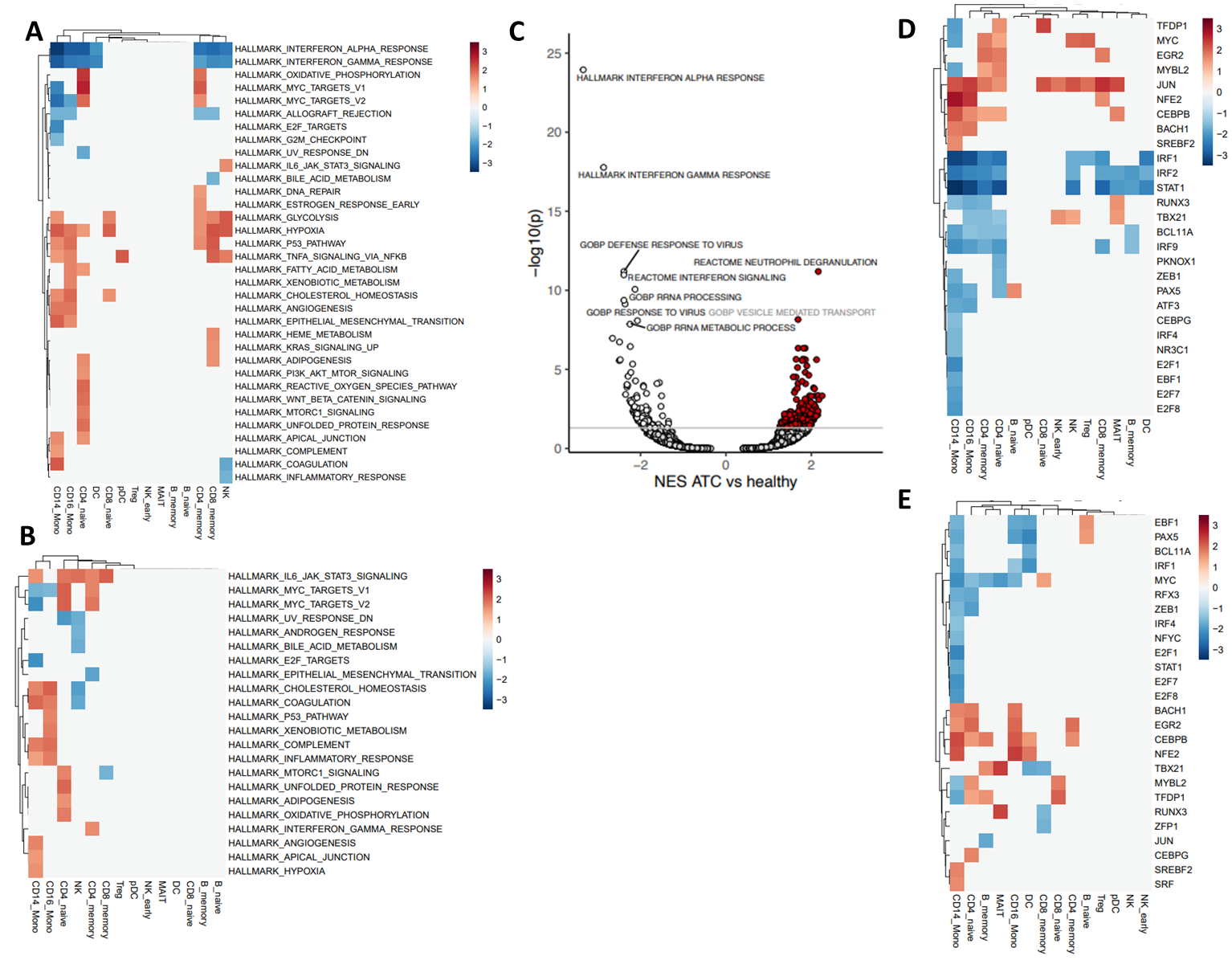

**Supplementary Figure 7**. Production of IL-6 **(A)**, IL-8 **(B)** and IL-1Ra **(C)** after 24 hours of stimulation or peripheral mononuclear cells with Pam3Cys or *Candida albicans*.

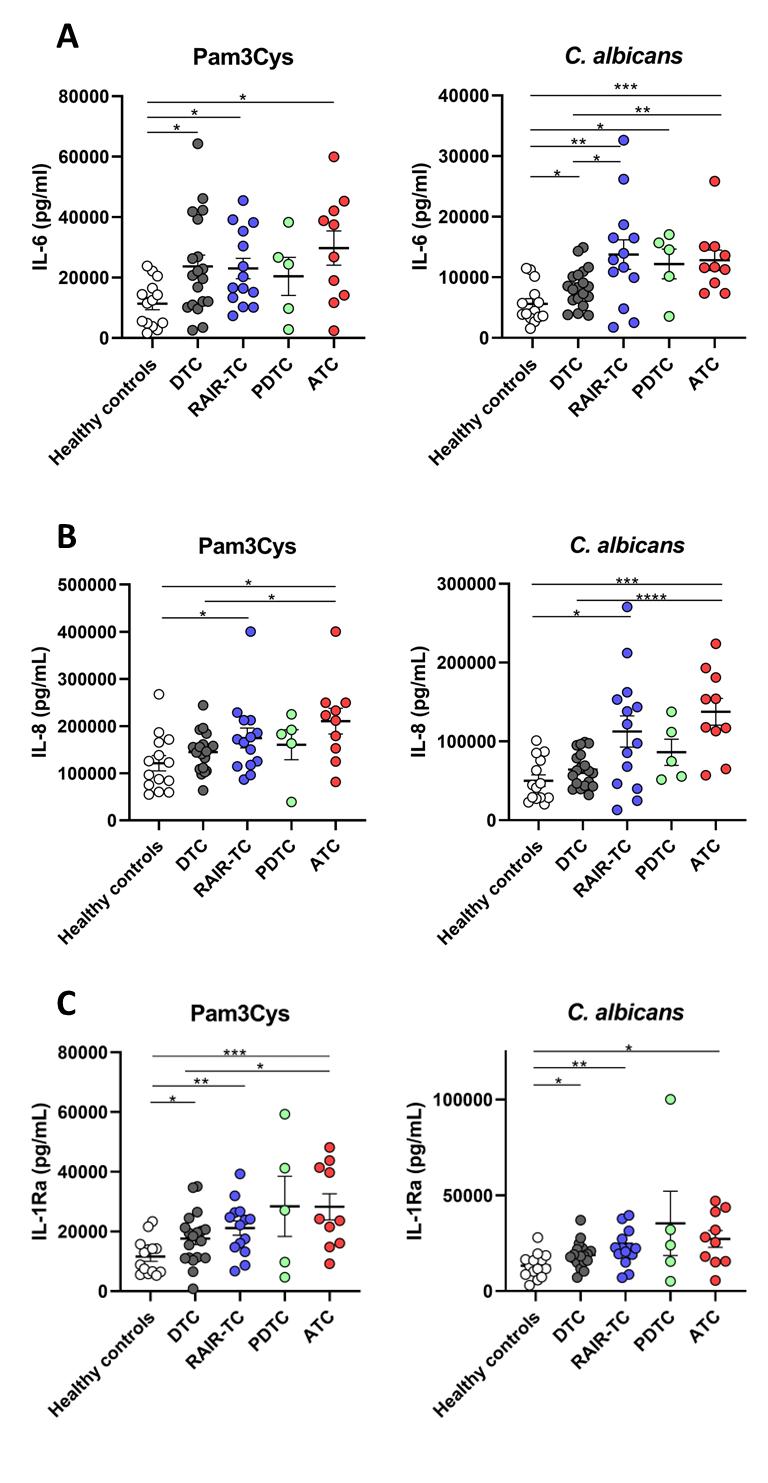

*: *p*<0.05 **: *p*<0.01 ***: *p*<0.001 ****: *p*<0.0001. DTC: differentiated thyroid carcinoma; RAIR-TC: radioiodine refractory thyroid carcinoma; PDTC: poorly differentiated thyroid carcinoma; ATC: anaplastic thyroid carcinoma.

**Supplementary Figure 8.** Production of IL-1β **(A)** and IL-10 **(B)** after 24 hours of stimulation of peripheral mononuclear cells by LPS, Pam3Cys, IL-1α or *Candida albicans.*

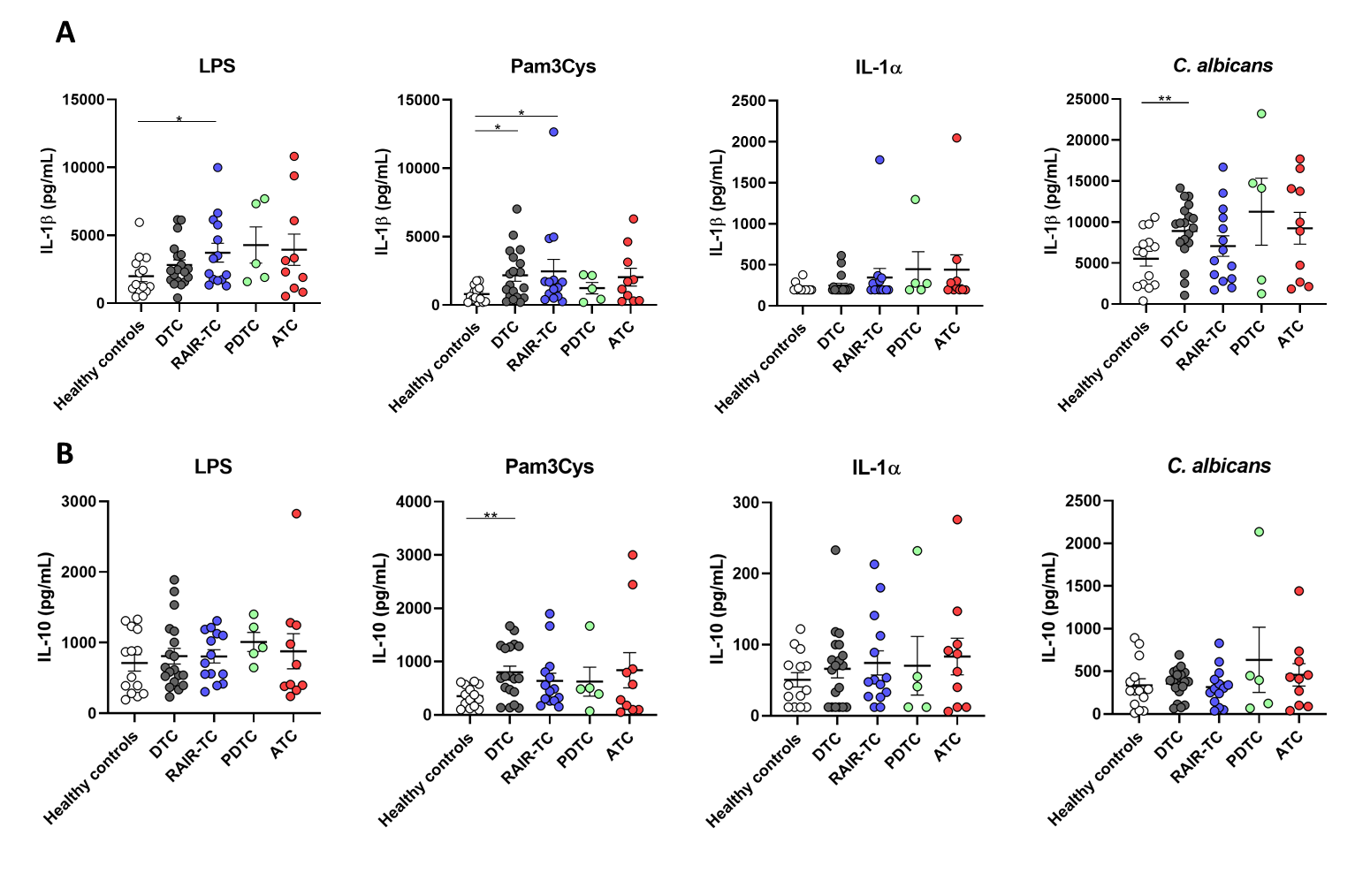

*: *p*<0.05 **: *p*<0.01 ***: *p*<0.001 ****: *p*<0.0001. DTC: differentiated thyroid carcinoma; RAIR-TC: radioiodine refractory thyroid carcinoma; PDTC: poorly differentiated thyroid carcinoma; ATC: anaplastic thyroid carcinoma.

**Supplementary Figure 9.** Production of IL-17 and IL-22 after 7 days of stimulation of peripheral mononuclear cells by *Candida albicans* or *Staphylococcus aureus*.

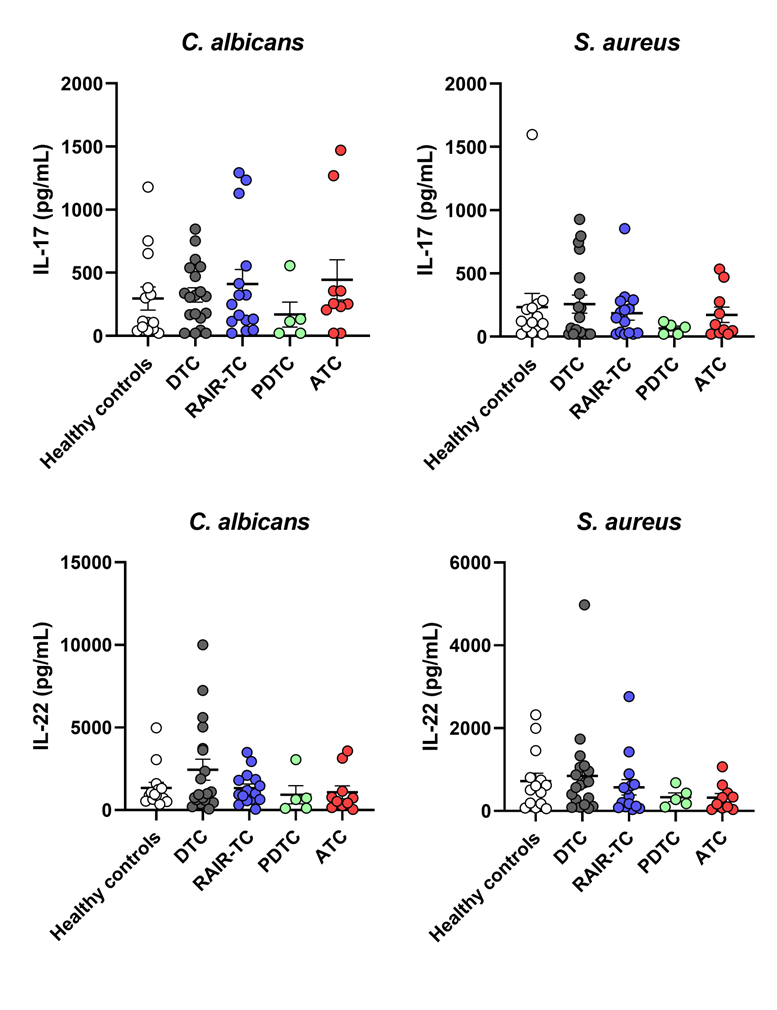

*: *p*<0.05 **: *p*<0.01 ***: *p*<0.001 ****: *p*<0.0001. DTC: differentiated thyroid carcinoma; RAIR-TC: radioiodine refractory thyroid carcinoma; PDTC: poorly differentiated thyroid carcinoma; ATC: anaplastic thyroid carcinoma.

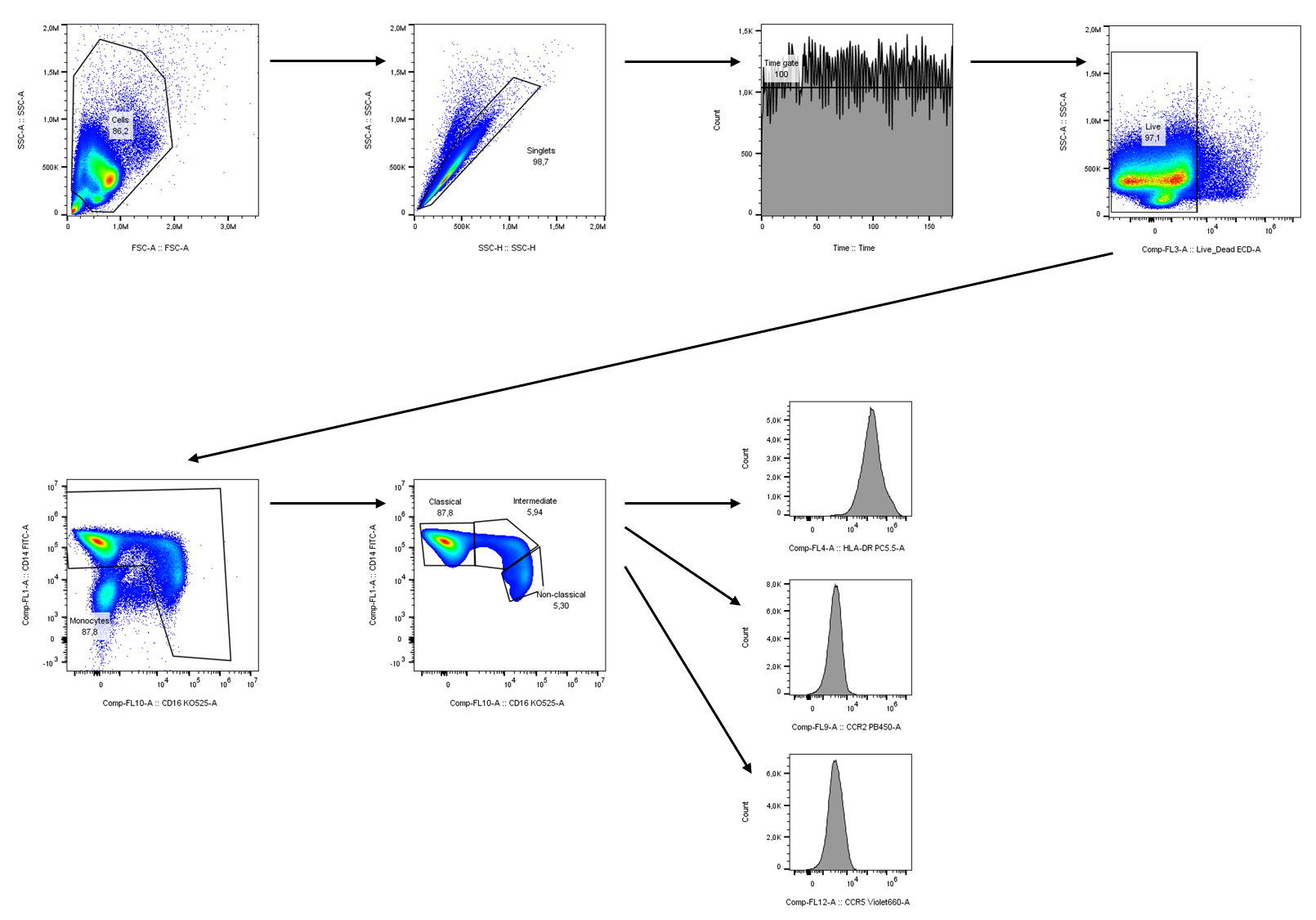
**Supplementary Figure 10.** Strategy for gating subpopulations of monocytes and and the expression of HLA-DR, CCR2 and CCR5.

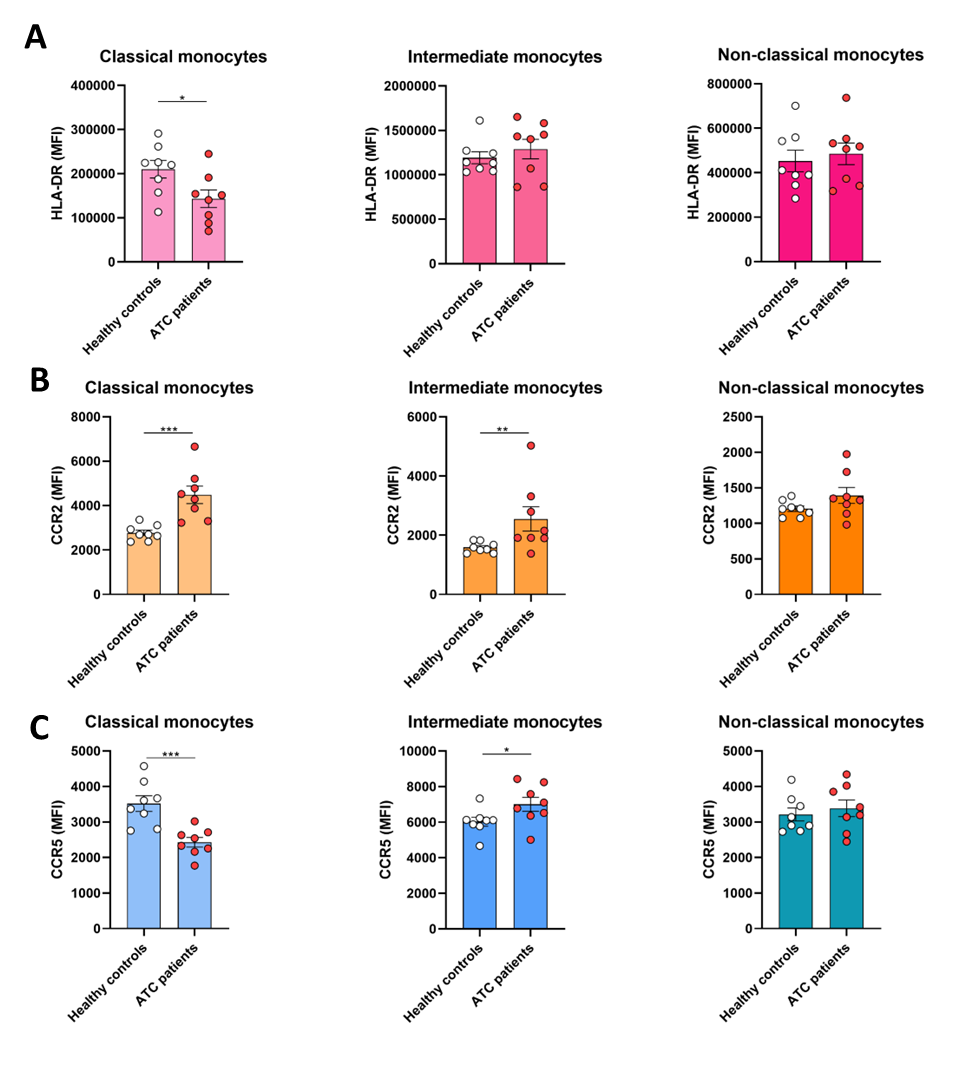
**Supplementary Figure 11.** Expression of markers HLA-DR **(A)**, CCR2 **(B)** and CCR5 **(C)** on subpopulations of circulating monocytes from healthy controls and ATC patients.

*: *p*<0.05 **: *p*<0.01 ***: *p*<0.001. ATC: anaplastic thyroid carcinoma.

**Supplementary Figure 12.** Viability of ATC cell lines 8505C, CAL62 and 8305C after 48 hours of treatment with different concentrations of IL-6 **(A)** or oncostatin M **(B)**.

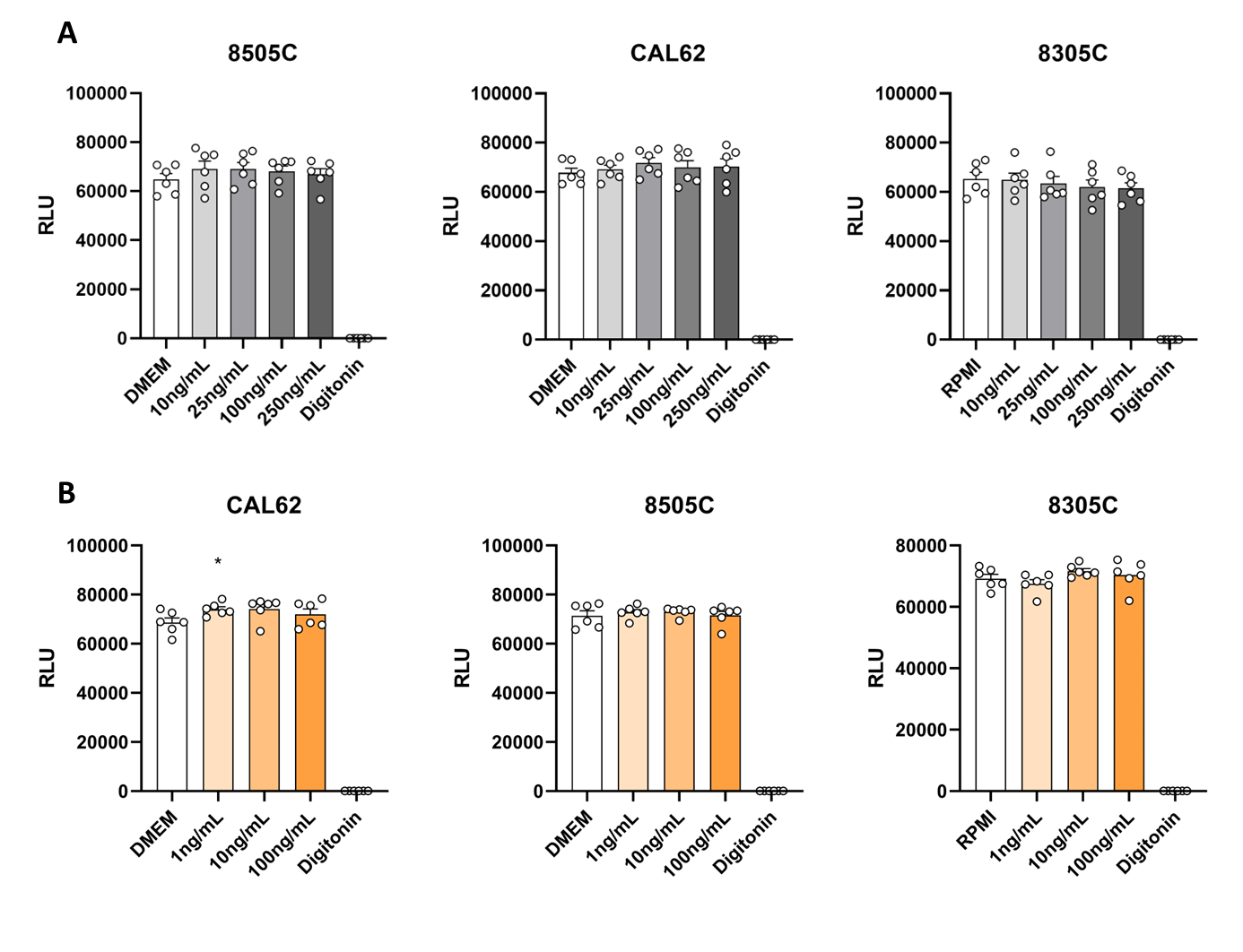

*: *p*<0.05. DMEM: Dulbecco’s Modified Eagle’s Medium. RLU: Relative light unit.

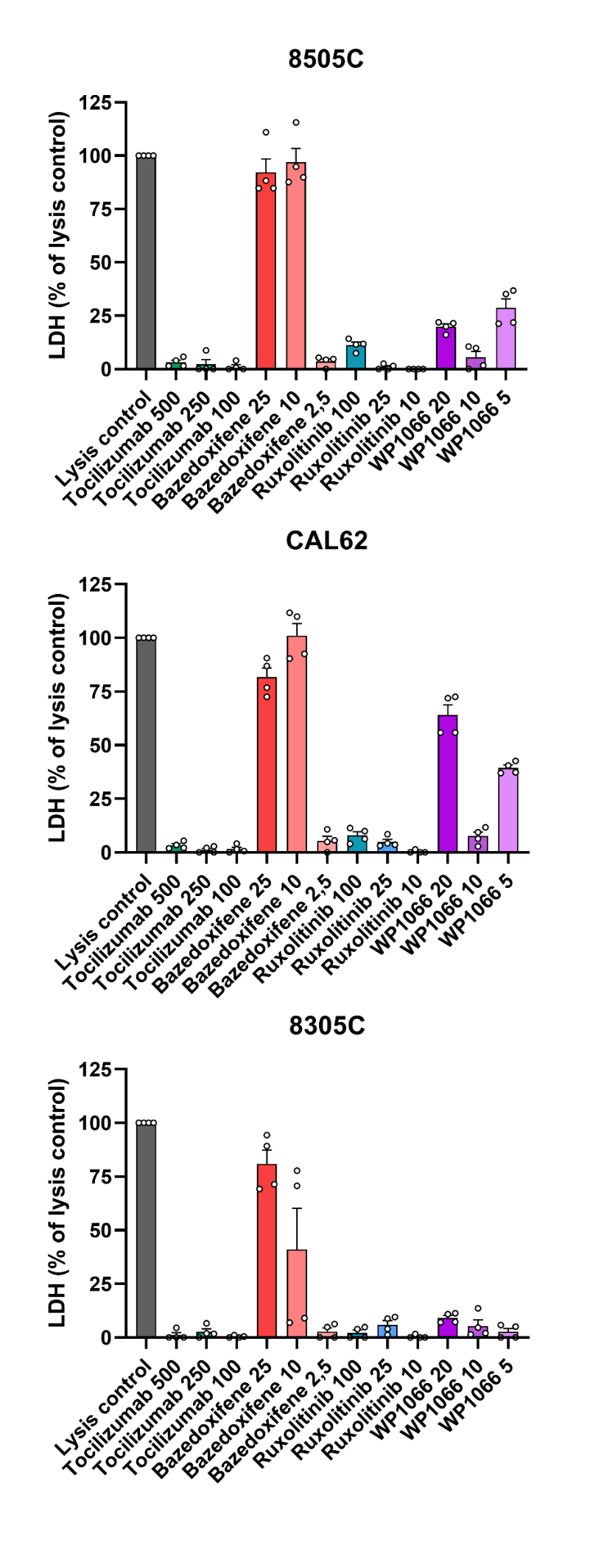
**Supplementary Figure 13.** LDH production by ATC cell lines 8505C, CAL62, and 8305C as percentage of lysis control after 48 hours of treatment with different concentrations of different compounds.

**Supplementary Methods**

*Materials*

*Escherichia coli* lipopolysaccharide (LPS; serotype O55:B5; Sigma-Aldrich) was repurified as previously described (51). Pam3CysK4 (P3C) was obtained from EMC Microcollections. Recombinant human interleukin-1α (IL-1α) was obtained from R&D Biotechne. *Candia albicans (C. albicans*; ATCC MYA-3573 (UC 820)) and *Staphylococcus aureus (S. aureus*; (Rosenbach ATCC 25923)) were grown overnight at 37°C in Sabouraud and Brain Heart Infusion broth, respectively. Microorganisms were harvested by centrifugation, washed twice in phosphate-buffered saline (PBS), and resuspended. *C. albicans* yeasts and *S. aureus* were heat-killed for 30 minutes at 95 °C.

*RNA single-cell sequencing analysis*

FASTQ data were processed using the cellranger multi pipeline. Pipeline outputs were used to generate a Seurat object for analysis in R using the Seurat package (1). Low complexity cells (log10(# detected genes/# detected UMIs) < 0.75), cells with high fraction of mitochondrial reads (> 7%) and cells with low number of detected genes (< 1000) were excluded from further analysis. scDblFinder() was run for doublet detection and cells with doublet scores over 0.25 excluded from analysis (2). Count data of the remaining cells were then further processed using NormalizeData(), FindVariableFeatures(nFeatures = 2000) and ScaleData(). Linear dimensional reduction was then performed using RunPCA(). The resulting PCA was batch corrected using the Harmony algorithm in IntergrateLayers() with each sample as a batch (3). The top 40 components of the resulting batch corrected dimensional reduction was used for clustering using FindNeighbors() and FindClusters() as well as nonlinear dimensional reduction using RunUMAP().

For celltype annotation, gene expression of established flow cytometry surface markers was examined in DotPlot() and FeaturePlot(). Clusters with highly similar marker expressions were combined and small clusters with ambiguous marker gene expression excluded from further analysis.

Differential gene expression of each identified celltype was analyzed in pseudobulk, to account for the donors as biological replicates in the statistical analysis. Pseudobulk profiles were generated as DESeq2 datasets using AggregateExpression() and DESeqDataSetFromMatrix(), comparing between healthy donors, ATC and DTC, including sex as a covariate (4).

For pathway analysis of differential expression results, genes were ranked by their DESeq test statistic (“stat” column) for gene set enrichment analysis (GSEA) using fgsea(). MSigDB Hallmarks, GO:BP, CP:KEGG_LEGACY and CP:REACTOME pathways were tested.

For use of the python implementation of SCENIC, the Seurat object was converted to an AnnData object using as.SingleCellExperiment() (SingleCellExperiment) and as_AnnData() (5). A transcription factor to gene adjacency matrix was generated using pyscenic grn. From this, regulons were pruned based on presence of a transcription factor motif in the hg38_10kbp_up_10kbp_down_full_tx_v10 cisTarget database using pyscenic ctx. The 255 identified regulons were then used as gene sets for pseudobulk GSEA.

*Bulk ATAC-sequencing*

Suspension cells were washed twice in ice cold PBS. The pellet was resuspended in 1:1 ice cold PBS and 2x lysis buffer (1 M Tris/HCl pH 7.5, 5 M NaCl, 0.5M MgCl2, 10% NP40). The cells were then centrifuged (300 x g) for 30 min at 4 °C and the pellet was resuspended with 24 µL clean-up buffer (5 M NaCl, 0.5 M EDTA, 10 mg/mL Proteinase K, 10% SDS) all the while kept on ice. Then, 1 µL Tn5 was added to each sample for tagmentation. The nuclei were heated for 6 min at 37 °C with 650 rpm agitation. Immediately after incubation, 9 µL clean-up buffer was added to each sample and incubated for 30 min at 40 °C with 650 rpm agitation. The samples were purified with a normal phase 2x SPRI purification and amplified by PCR using Nextera primers (Illumina) and KAPA HiFi HotStart ReadyMix (Roche). Tagmented DNA was amplified using a 5 cycle PCR protocol. A reverse phase 0.65x SPRI bead purification step was performed, followed by a 1.5x SPRI beads clean-up. Next, a second PCR amplification program was used to further amplify the tagmented DNA. The program was identical to the first PCR program, including the Nextera primers, except for the number of cycles, which was determined by qPCR. Real-time amplification was performed using iQ SYBR Green mix (Bio-Rad) on a CFX96™ Real-Time System (Bio-Rad) and quantified using Bio-Rad CFX Manager. The Ct value + 2 was used as PCR cycles for the second PCR amplification step. Then, two successive 1.5x SPRI clean-ups were performed to generate the ATAC libraries. The DNA was stored at -20 °C and an aliquot was used to measure the DNA concentration with a Qubit® dsDNA HS Assay Kits (Life Technologies). Library size distribution was measured using High Sensitivity DNA analysis (Agilent) on an Agilent 2100 Bioanalyzer and its corresponding software. DNA fragment distribution followed a mononucleosomal-like pattern at approximately 200 bp and 320 bp. The libraries were sequenced using a NextSeq 2000 system (Illumina). Reads were trimmed, aligned, and counts were generated using the seq2science pipeline with default settings (6). Peaks were called using MACS2 (7) and differentially accessible regions were determined using DESeq2 (8). Motif enrichment analysis was performed using HOMER (<http://homer.ucsd.edu/homer/motif/>), using peaks with adjusted P-values < 0.05.

*Cytokine and lactate assays*

IL-1β, IL-6, IL-8, IL-10 and IL-1Ra were measured in supernatants of 24 hours stimulated PBMCs and IL-17, IL-22 and (IFN-γ) in supernatants of 7 days stimulated PBMCs by commercial ELISA kits according to the instruction of the manufacturer (R&D Biotechne). Lactate concentration was measured in supernatants from 24 hours stimulations using a lactate oxidase based enzymatic reaction (Sigma). H_2_O_2_ from the reaction was coupled to the conversion of Amplex Red reagent (Life Technologies) by horseradish peroxidase (Sigma). The resulting fluorescence (at 530/25 and 590/35 nm) was then measured immediately.

Circulating concentrations of IL-6, IL-1Ra, granulocyte-colony stimulating factor (G-CSF), granulocyte-macrophage colony-stimulating factor (GM-CSF), osteopontin and chemokine (C-X-C motif) ligand 9 (CXCL9) were measured in EDTA plasma using Quantikine ELISA kits (R&D). Circulating IL-1β concentration was measuring in EDTA plasma using a Simple Plex cartridge on an ELLA automated immunoassay system (R&D Biotechne).

*Analysis of proteomics*

Analysis of the Olink proteome assay was performed in R version 4.1.3. Proteins that were detectable in <75% of the samples were excluded from the analyses. Normalized Protein Expression (NPX) values are expressed on a log2 scale. A principal component analysis (PCA) was used to compare the protein expression pattern between all subgroups. NPX values of proteins were compared between subgroups by the Mann-Whitney U test. Package “ggplot2” was used for visualization. False discovery rate (FDR) corrected *p*-values were used to adjust for multiple testing.
